# Potential benefit of loss-of-function on bacterial fitness

**DOI:** 10.64898/2026.08.13.744710

**Authors:** David Hidalgo, Lizeth Soto-Ávila, O. Alejandro Aguilar-Vera, Daniela Ledezma-Tejeida, José A. Farías-Rico, José Utrilla

## Abstract

Escherichia coli is a well-studied organism with extensive genomic and proteomic data. This study examines how gene loss reallocates cellular resources and impacts fitness. Genes were classified based on fitness measurements as essential, important, mean-effect, or fitness-enhancing. Using proteomic data, we analyzed the relationship between protein production cost and fitness, finding that genes with a high proteomic mass fraction are more likely to affect fitness, while fitness-enhancing deletions rarely improve fitness by reducing proteomic burden. We calculated the cumulative of proteome fractions encoded by genes classified as mean-effect and compared it with the results from the ME-model simulations. The mean-effect category constitutes 31-75% of the proteome, with the highest proportion LB, while enrichment analysis of core mean-effect genes highlighted transmembrane transport as the main functional category. Furthermore, we identified a subset of genes whose deletion increased fitness compared to the mean; they generally have low expression, and many have unknown functions. AI-assisted structural analyses identified domains and conserved features compatible with DNA-binding proteins, suggesting that some may represent putative transcriptional regulators requiring further validation. RpoS, stress sigma factor controlling up to 15% of the proteome is one of the transcriptional regulators in the fitness-enhancing category. Our findings suggest that the cost of being a generalist is linked to transcriptional regulation, while molecular transport represents a high burden for nutrient readiness.

**Importance:** This study provides new insights into how gene loss benefits bacteria by identifying gene categories and their associated protein fractions whose disruption does not impose large fitness penalties. Additionally, it uncovers specific fitness-enhancing genes and generates hypotheses based on structural analyses for previously uncharacterized ones. Our findings suggest that several of these genes may encode putative transcriptional regulators, highlighting a potential role for regulatory complexity in cellular efficiency. By revealing how certain gene deletions enhance fitness and which gene categories are nonessential, this work advances our understanding of bacterial adaptation and genome streamlining. These insights have broad implications for evolutionary biology, metabolic engineering, and biotechnology, offering strategies to optimize microbial function by selectively reducing genetic and regulatory burden.

## Introduction

Generalist organisms like *Escherichia coli* are able to thrive across diverse environments, evolving the ability to adapt to naturally fluctuating conditions through the expression of bet-hedging functions(Zhu and Dai, 2024). Yet, this adaptability involves a trade-off, incurring in costs that often curb cellular growth. Intriguingly, gene loss, as documented in evolutionary contexts (Albalat and Cañestro, 2016), emerges as a pervasive mechanism allowing organisms to optimize their genetic repertoire when particular genes are unnecessary in certain environments.

Even in the extensively studied model bacterium *E. coli*, approximately 50% of its genes are not well-characterized, with up to 35% lacking experimental evidence behind their function(Moore et al., 2024). Traditionally, genes are categorized as essential or non-essential based on their deletion outcomes under standard conditions, such as growing in a rich medium. However, this binary classification under a single condition fails to capture biological intricacies that define the function of each gene. The emergence of barcoded single-gene mutant libraries (Wetmore et al., 2015) presents an unparalleled opportunity to explore *E. coli* single gene mutant physiology and gene function on a coarse quantitative level.

By reducing the expression of superfluous genes, a process managed through the interplay of sigma factors and transcription factors within the Transcriptional Regulatory Network (TRN), organisms can reallocate resources efficiently(Fang et al., 2017). This enhances their overall fitness and adaptability under specific environmental conditions. Mutations in genes involved in central processes, like those encoding for the RNA Polymerase (RNAP), can reprogram a cell’s regulatory network to reduce the expression of unnecessary functions. These mutations frequently arise in adaptive laboratory evolution experiments. Studies have showcased the fitness trade-offs these mutations entail (Utrilla et al., 2016). Genome-scale models of metabolism and gene expression (ME-models) provide insights into how protein utilization varies across environments. Moreover, comparing protein abundance in these contexts reveal many functions are unnecessarily expressed. These findings substantiate that the costs inherent to being a generalist organism are quantifiable, highlighting the need to investigate potential fitness gains from losing specific functions.

Here, we employed a functional genomic approach, integrating fitness datasets, absolute proteomics data, ME-model simulations, and functional gene categorization to deliver a quantitative cellular economic assessment of the potential benefits of loss of function across environments in *E. coli*. Our results reveal that most genes do not largely contribute to fitness in fixed environments, we investigate functional categories associated with burden and show that gene expression regulation in static environments may be suboptimal with selection primarily driven by changing conditions. We show that our findings can be generalized to other bacteria. By extending our analytical framework to include mutational and gene expression data from the Lenski Long-Term Evolutionary Experiment (LTEE) and other divergent species.

## Methods

### Fitness data

Previously reported fitness data (Price et al., 2018) for *E. coli* and other bacteria was used for this study. A fitness assay reports the log2 of final/initial abundances of randomly barcoded transposon mutants (RB-TnSeq). A total of 46 standard conditions with duplicate measurements were considered for this study. In every individual experimental condition, genes were classified according to their measured fitness value when disrupted as: a) between -0.5 to 0.5: mean-effect, b) less than -0.5 and higher than -2: important, c) lower than -2: essential, and d) greater than 0.5: fitness-enhancing. Genes with no data were considered essential and classified as such.

### Note on Definition and Interpretation of Fitness Effects

Fitness scores for each gene knockout were calculated from barcode frequency changes. We categorized genes by comparing scores to the central peak (mode) of the distribution in each condition. Genes within this peak are classified as “mean-effect”. This mode-based framework is deliberate. Fitness distributions in pooled transposon assays are strongly left-skewed by deleterious knockouts, making the arithmetic mean an unreliable proxy for biological neutrality. The term mean-effect therefore reflects that these knockouts exhibit fitness changes near the center of the observed distribution, rather than implying neutrality relative to wild-type. Importantly, this classification does not assume that mean-effect knockouts have zero fitness cost relative to wild-type, but rather that any effects are subtle and not resolvable given the structure and resolution of pooled barcode frequency data in a given environment. To place these results in broader context, we compared pooled TnSeq fitness values with relative growth measurements from the Tong single-deletion dataset (Tong et al., 2020) (Supplementary Figure S7). As expected, strong fitness defects are concordant between datasets, while genes classified as mean-effect show broad agreement at a coarse level, consistent with the known weak correlation between competitive fitness and colony growth for small-effect mutations.

### Fitness-proteomics correlation and analysis

A quantitative *E. coli* proteomics dataset (Schmidt et al., 2016) containing protein mass (in femtograms, fg) and protein copies per cell on several growth conditions was used. A total of eleven conditions were selected for this study: LB medium and M9 minimal medium with either Glucose, Acetate, Glucosamine, Glycerol, Pyruvate, Xylose, Mannose, Galactose, Succinate or Fructose as the carbon source.

The eleven experimental conditions are shared between the proteomics and fitness datasets. For each condition, protein mass distributions were compared between genes classified as mean-effect and genes in each other fitness categories using the non-parametric Kruskal-Wallis test, as the data were not normally distributed. The resulting P-values were adjusted for multiple testing using the Holm-Bonferroni method.

### Proteomic mass fraction calculations

To determine the proteomic mass fraction of GO terms we considered the sum of the proteomic loads of the genes on the GO term in a specific condition. For the calculations on RpoS targets we compared the proteomic mass fraction of the total set of genes regulated by RpoS (sigma 38), regardless of whether it is co-activated by another sigma, with the set of genes exclusive to RpoS. Since the proteomic load varies according to the growth condition, we show the proteomic mass fraction as a percentage of the theoretical fraction of the proteome released per condition.

### Fitness-enhancing gene determination

From the fitness dataset and considering all 46 conditions mentioned above, we determined the number of conditions in which each gene falls into the “fitness-enhancing” category. From there, we ordered the genes that more frequently appear as fitness-enhancing (Supplementary Table S19).

### Prediction of function of y-genes

We employed a combination of two approaches to infer the functions of the target proteins: sequence-based comparisons and structure-based analysis. Sequence based annotation was performed by Hidden Markov Model comparisons employing the HH-suite3 software for fast remote homology detection and deep protein annotation (Steinegger et al., 2019) (38) implemented in the MPI toolkit web-server (Gabler et al., 2020). We built HMM profiles for each protein target and we compared them with a variety of HMM databases. We recorded alignments of the target proteins and their hits. Probability is the most important score in HMM comparisons. If the HMM profiles from two proteins are aligned with more than 90% probability their homology is almost certain. Also, similarities among the top targets were used to strengthen the inference of homology.

For structural analysis models were generated with alpha fold 2 (Jumper et al., 2021) implemented in collabfold (Mirdita et al., 2022) and ESMfold (Lin et al., 2023). Once the models were generated, we performed structural searches employing DALI (Holm et al., 2023). Detailed structural alignments were performed with PDBefold (Krissinel and Henrick, 2004). Z-scores were recorded. Typically, a Z-score higher than 20 strongly indicates that two structures are homologous. A score between 8 and 20 suggests probable homology, while scores between 3 and 8 fall into an uncertain grey area. A Z-score below 3 is generally considered insignificant. If two structures were considered structural homologues according to the above indicated scores, we assumed they performed similar functions. Full data with scores is included in supplementary Table S17 (High confidence) and supplementary Table S18 (Low confidence).

### ME-model simulations

To determine the translation fluxes, we simulated cellular growth using the *Escherichia coli* iJL1678b-ME genome-scale-model(Liu et al., 2014). This model quantitatively accounts for both metabolism and gene expression. In order to improve our predictions, we first updated the effective reaction rate constants (k_effs_) using a validated kinetic profiling method(Heckmann et al., 2018). This modification integrates protein abundances using the same quantitative proteomic data mentioned above to better reflect physiological conditions. All model manipulations were made using Python 3.6.13 (Van Rossum and Drake, 2011) and the COBRAme library (Lloyd et al., 2018). The model was solved using the qMINOS solver(Ma et al., 2017).

### Functional Gene Ontology (GO) Enrichment Analysis

All fitness category genes were annotated using GO annotations from Bioconductor (Gentleman et al., 2004) for Biological Process. Functional enrichment tests were performed using the ViSEAGO R package (Brionne et al., 2019). A functional enrichment, supported by GO terms, within each fitness category of genes from *E. coli* was determined using Fisher’s exact tests and the “elim” algorithm (p < 0.01), with all of the expressed genes used as the background(Alexa et al., 2006). GO enrichment analyses were performed independently for each growth condition. The complete lists of enriched GO terms are provided in Supplementary Tables S2–S12, corresponding to glucose, LB, glucosamine, glycerol, xylose, mannose, galactose, fructose, acetate, pyruvate, and succinate, respectively. We then summarized enriched GO biological processes by growth condition and fitness category. For each enriched GO term, the adjusted proteome fraction was calculated by summing the protein-level proteome contributions assigned to that term within each fitness category. The resulting GO-level summary is reported in Supplementary Table S13.

Genes categorized as fitness-enhancing in more than 10 conditions were extracted, and Transcription Factors were identified based on their common name. Transcription factor-gene regulatory interactions were retrieved from the RegulonDB database(Salgado et al., 2024). A gene with regulatory redundancy was defined as any gene having two or more Transcription Factors having the same effect (activation/repression) over the gene. Analysis and statistical tests were performed with custom scripts in the R software.

## Results

Our analysis leverages fitness scores obtained from previously reported single-gene mutant libraries generated by random bar coded transposon sequencing (RB-Tnseq) (Price et al., 2018). This dataset covers 3789 of the ∼4600 genes in *E. coli*, accounting for 84% of the genome. Our primary goal was first to assess the distribution of fitness values across 46 different growth environments (defined here as distinct compositions of growth media), categorize genes based on their contribution to fitness in each environment and across multiple environments, and determine their associated protein mass.

Fitness scores are numerical measures of a mutant strain’s reproductive success in a defined growth condition, which is quantified by tracking its barcode’s abundance. It is calculated as the base-2 logarithm of the ratio of final to initial abundance of each barcode-tagged mutant, which indicates how much its frequency has changed over a few generations. Values close to zero represent gene disruptions with fitness effects near the mean of the distribution, reflecting subtle effects that cannot be interpreted as strictly neutral. Positive values indicate a growth advantage compared to the mean, and negative values a disadvantage. Most genes have fitness values near zero, meaning that their disruption has a marginal average effect on fitness, often referred to as “no-phenotype” (Price et al., 2016, 2018). Values can range from -10 (gene is essential) to 10 (disruption is beneficial). We used the fitness scores to classify mutants in several categories: mean effect (-0.5 to 0.5), Important (-2 to -0.5), Essential (below -2), and Enhancing (above 0.5) (see methods for a more detailed note on the interpretation of fitness scores). This reflects the assay’s dynamic range, where |fitness| < 1 is subtle and |fitness| ≥ 2 is strong. Genes without fitness data (e.g., from no insertions or low read counts) were grouped with Essential genes, since it is likely the absence of the mutant implies a deleterious mutation. Fig. 1A shows a typical fitness score distribution of the single gene mutants in a medium with glucose as the only carbon source.

**Figure 1.**
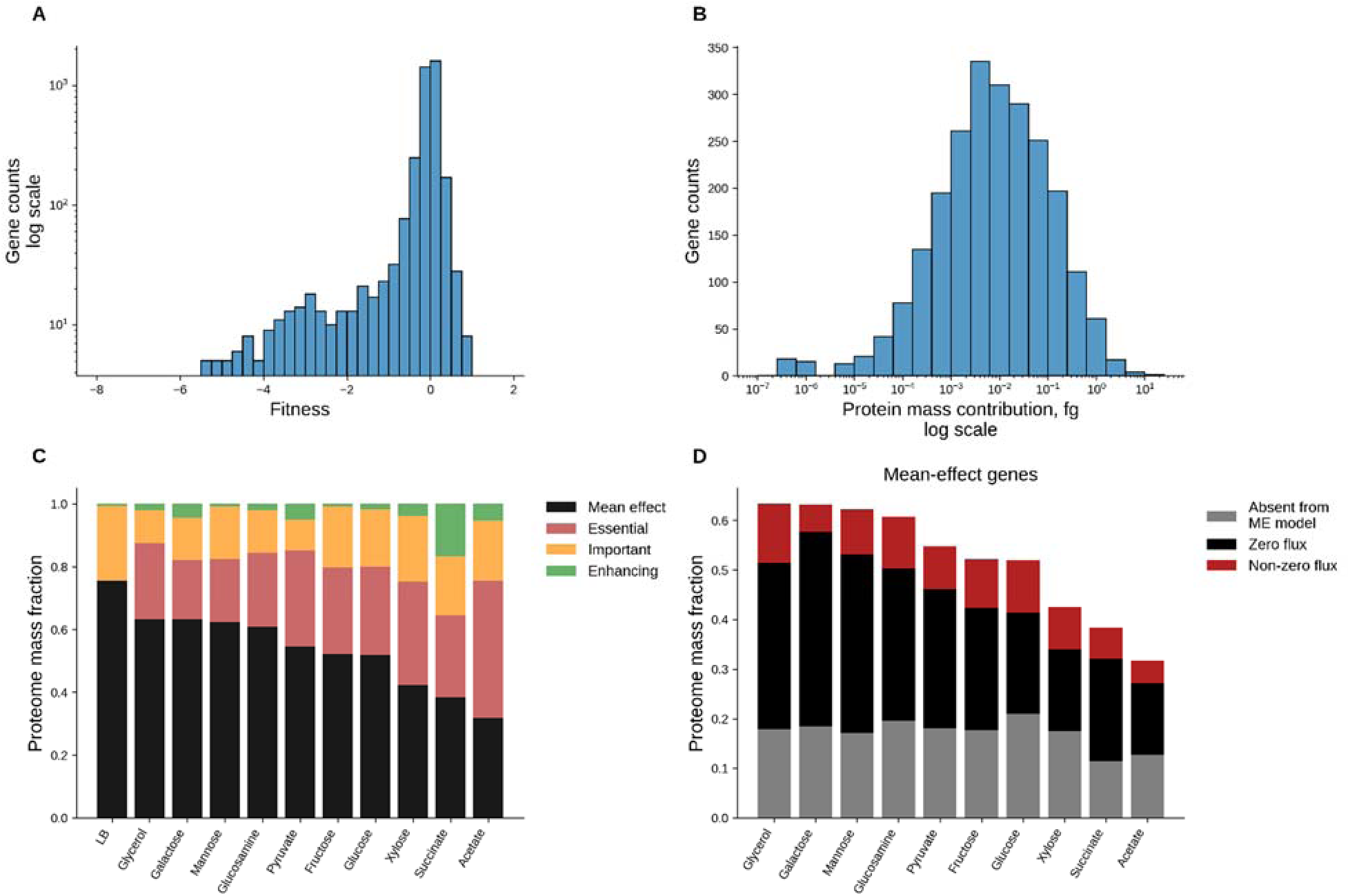
Fitness as a function of protein cost. A, B Typical distributions of fitness and protein mass contribution respectively. Data from glucose assays was selected for the histograms. All protein mass and fitness data from individual conditions follow a non-normal distribution per a Shapiro-Wilk test with *p* = 0.001. C, Proteome mass fractions of each fitness category across the eleven shared conditions of fitness and proteomics. D, Proteome mass fractions of mean-effect genes and their classification as either absent from the ME-model, present in the ME-model but not used in the solution (zero flux) or present in the ME-model with predicted flux in the solution across conditions.

As for the protein mass, we examined a thorough quantitative proteomics dataset (Schmidt et al., 2016) that includes the mass contribution to the proteome (i.e., a protein’s mass in the cell, not its molecular weight) of 2289 proteins measured individually for each of the 22 experimental conditions. The mass contribution of a protein per cell ranges from 10^−7^ to approximately 34 femtograms (fg), with a median value around 10^−3^ fg (Fig. 1B). In every condition, about 7-11% of proteins account for 80% of the total measured proteome mass. There is an overlap of 1951 genes and eleven experimental conditions between the fitness and proteomics datasets using the same strain: *E. coli* BW25113, allowing us to directly compare these two sets of data in these eleven conditions (Supplementary Table S1 integrates fitness and proteomic data).

### The mean-effect category constitutes up to 75% of the proteome

We previously showed that the unused part of the proteome may constitute up to 30% in glucose minimal medium(O’Brien et al., 2016; Lastiri-Pancardo et al., 2020). However, this unused fraction was calculated by the sum of all the proteins that are expressed in that condition, and do not have any contribution to the predicted proteome on a genome scale model of metabolism and gene expression (ME - model) (Lloyd et al., 2018). Here, we took a different approach, first we calculated the mass fraction of the total of proteins in each fitness category (Fig. 1C). Then, we compared the abundance of the proteins with mean-effect in fitness to the predicted use of those proteins by the ME-model, simulating batch growth in minimal media with each of the 10 measured carbon sources (Fig. 1D).

The total mass fraction of all the proteins in their corresponding fitness category displays contrasting behaviors: a) essential proteins generally do not contribute to the largest fraction of proteomic mass; an exception of this occurs in acetate, a very poor carbon source for *E. coli*. The proteins from essential genes account for less than half of the proteome mass (median = 0.26, fractions go from 0.43 in acetate down to 0.001 in LB). b) The mean-effect category is the largest fraction contributing to the proteome, going from 0.75 down to 0.31 (median = 0.54) depending on the growth condition. c) The important (median = 18) and enhancing (median = 0.02) categories of proteome represent the smallest fractions (0.24 to 0.34). We further analyzed the proteins from the mean-effect genes using the ME-model solution for each independent condition (Fig. 2B). We found that a predominantly constant fraction of ∼0.2 of the proteome is not included in the ME-model, however most of the mean-effect proteins are included in the ME-model reconstruction but are not predicted to be expressed in that specific condition. Furthermore, a portion of the mean-effect proteins is predicted to be expressed as they show a translation flux in the ME-model simulations (Fig. 1D). We excluded LB medium from the analysis due to its undefined chemical composition. Unlike defined media, the variable and complex nutrients in LB make it unfeasible to accurately model the consumption rates of its individual components without introducing arbitrary adjustments to influx reaction rates. Overall, this analysis presents the use case of an ME-model to be a guide to calculate the amount of proteome that could be spared if the mean-effect protein expression could be eliminated, with fractions ranging from 0.51 to 0.27.

**Figure 2.**
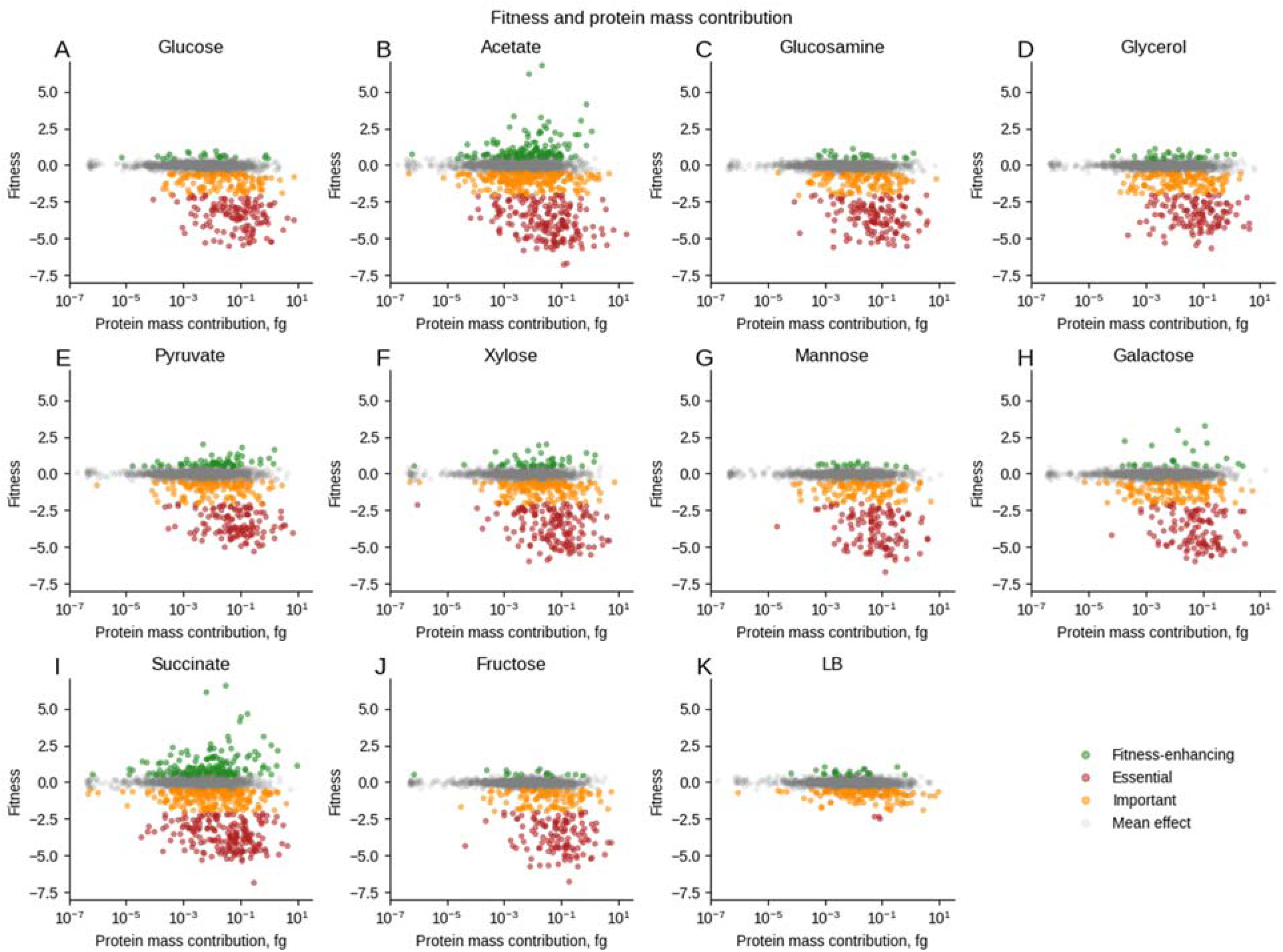
A-K, Correlation between fitness and protein mass. Single RB-TnSeq mutant fitness values are plotted as a function of protein weight in the eleven shared conditions between fitness and quantitative proteomics datasets. Colors indicate fitness classification: 1) gray, mean-effect, 2) orange, important, 3) red, essential, and 4) green, fitness-enhancing.

### Heavier proteins are more likely to impact the fitness

As previously noted, most genes had fitness values close to zero across all conditions (Fig. 2 A-K, gray dots), with protein mass spanning several orders of magnitude. In all conditions except for LB, a subset of data deviated from zero on the fitness axis, typically skewing towards higher mass (fg) values. We tested whether each fitness category contained a distinct population of proteins based on protein mass. For each growth condition, we compared the mass contributions of the proteins in the mean-effect group with those of proteins belonging to each remaining category using a non-parametric Kruskal–Wallis test, followed by Holm–Bonferroni correction for multiple comparisons. This analysis showed that essential and important genes represented distinct protein-mass distributions from the mean-effect group in most conditions, achieving a statistical confidence level of predominantly 99.9% (adjusted p < 0.001; Supplementary Figure S1). A distribution analysis demonstrated an increase in median values for gene subsets different from the mean-effect category, reflecting their higher mass (fg) values. Notably, genes with smaller mass (fg) values (10^−7^ to 10^−4^ fg) generally fell outside these subsets. In six conditions, glucose, glucosamine, glycerol, mannose, fructose, and LB medium, fitness-enhancing genes did not represent a distinct population from the mean-effect group after Holm–Bonferroni correction at α = 0.05. In contrast, in acetate, pyruvate, xylose, galactose, and succinate, the fitness-enhancing category contained proteins that represented a distinct subset, with significance ranging from p < 0.05 to p < 0.001 depending on the condition. Across conditions, genes with higher mass (fg) values mostly belonged to the essential and important categories. Therefore, larger proteome contributions were associated with stronger, often negative, fitness effects. Across conditions, genes with higher mass (fg) values mostly belong to the essential and important categories. Therefore, losing genes encoding heavier proteins tends to produce a more substantial and often negative impact on fitness. Analysis of both the fitness and proteomics datasets revealed that genes encoding heavier proteins generally contributed more significantly to fitness outcomes than those encoding lighter proteins.

### Main enriched functional categories across carbon sources

To compare the functional composition of genes within each fitness category (essential, important, mean-effect, and enhancing), we performed Gene Ontology (GO) enrichment analysis (Ashburner et al., 2000) using annotations from the *org.EcK12.eg.db* R package, specific for *Escherichia coli* K-12. Enrichment was conducted with ViSEAGO, which implements topGO (Alexa et al., 2006) and its “elim” algorithm (developed in 2006) to account for the GO hierarchy and highlight biologically relevant terms.

Across the eleven tested carbon sources and the four fitness categories, we identified a total of 239 enriched GO terms, with an average of 118 enriched terms per condition. To focus on the most robust processes, we restricted the visualization to GO terms consistently enriched across all conditions. As shown in Fig. 3, these conserved enriched terms were detected in the essential, important, and mean-effect categories. The fitness-enhancing category was excluded, as it contained too few genes to identify GO terms enriched across all conditions.

**Figure 3.**
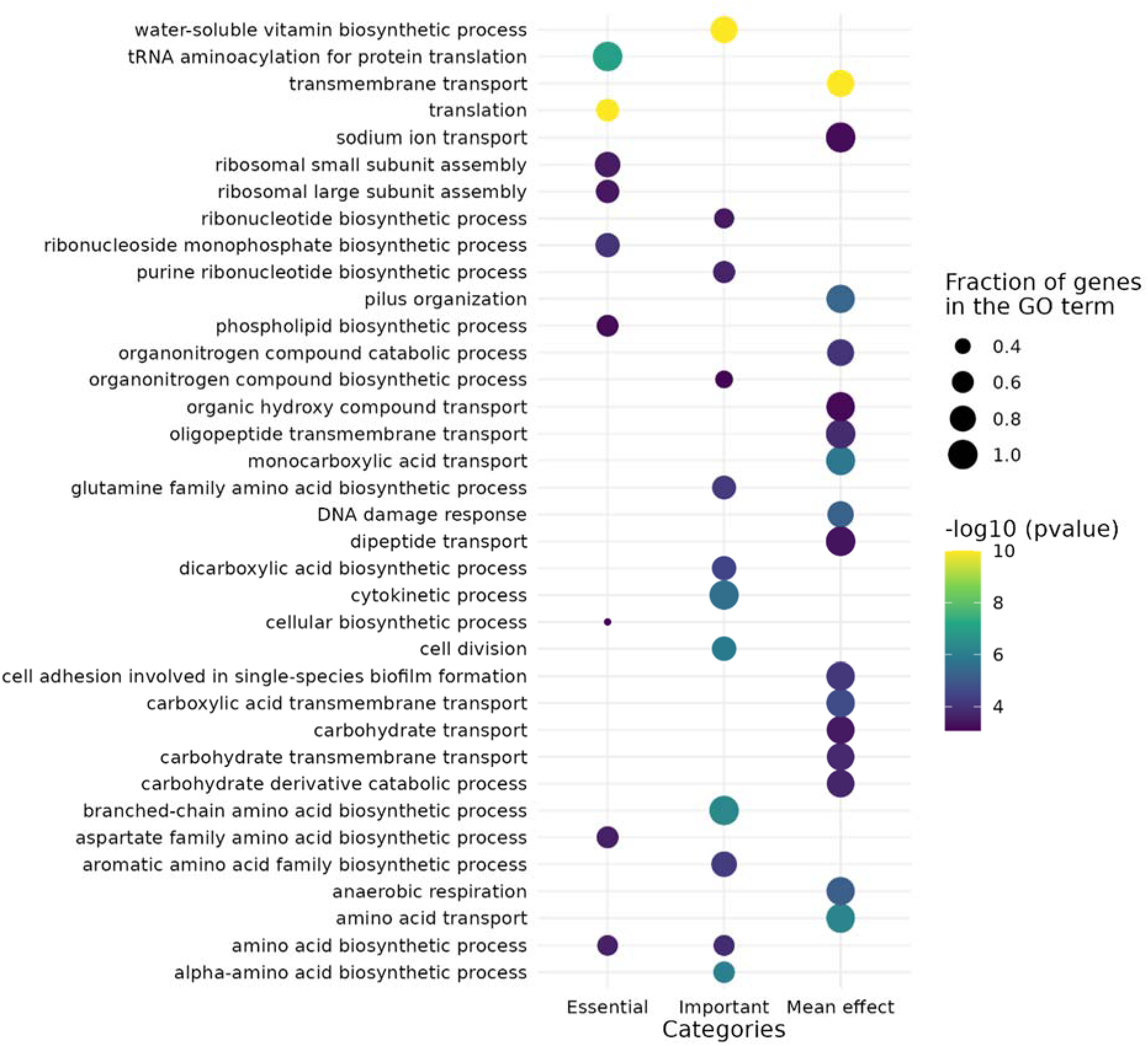
Dot plot of enriched Gene Ontology (GO) terms among the fitness categories across all different carbon sources. The size of the dot represents the fraction of genes in the GO term, and the color indicates the normalized enrichment of each term, determined by the *p*-value on a log scale (yellow is the most highly significant). A filter was applied for processes with a frequency greater than 0.25 and a *p*-value < 0.001. Note that the “enhancing” category did not present any shared process among the different carbon sources

We found that the essential and important categories are enriched in core biological processes such as translation, biomolecule biosynthetic processes and cell division. These results are consistent with previous reports, reflecting that these functions carry out key cellular roles. In contrast, the mean-effect category showed enrichment in processes related to the transport of various molecules, pilus formation, DNA repair, cell adhesion, and anaerobic respiration. These patterns suggest that genes with little apparent fitness effect are not functionally random, but are enriched in processes related to environmental interaction and nutrient use.

Having identified the enriched biological processes associated with each fitness category, we next asked whether these functions also represented a substantial proteomic investment. We first examined glucose minimal medium, the reference condition for the ME-model and the adaptive laboratory evolution experiments analyzed below. In glucose, we found a total of 117 enriched GO terms (Table 1). The essential genes covered the largest number of enriched functions (53 of 117 GO terms), the mean-effect category, which is the second most abundant, covers 45 GO terms, many associated with transport processes. To assess whether these functional patterns extended to another carbon source, we next applied the proteomic contribution analysis to galactose. Of the 117 GO terms identified under glucose, only 4 were shared between the essential and important fitness categories, showing a high specificity of the functions in this condition (Supplementary Table S2). The complete GO enrichment results for each individual growth condition are provided in Supplementary Tables S2-S12, whereas the corresponding adjusted proteome fraction per GO term and fitness category is summarized in Supplementary Table S13.

**Table 1.** Breakdown of gene and gene ontology (GO) term groupings for each fitness category under glucose condition. Column 1 shows the fitness category. Column 2 indicates the total number of genes identified within each category. Column 3 shows the number of enriched GO terms identified within each category. Column 4 reports the number of genes represented within the enriched GO terms. Column 5 indicates the number of proteins associated with proteomic data. Column 6 shows the summed proteome mass percentage across all enriched processes within each fitness category. Additionally, *four GO processes are shared across these categories.

| Fitness category | Total genes | Number of GO term enriched | Genes within GO term enriched | Proteins with data | Proteome mass (%) across all GO processes |
| --- | --- | --- | --- | --- | --- |
| Essential | 443 | 53* | 368 | 399 | 40.3 |
| Important | 209 | 20* | 130 | 128 | 7.12 |
| mean-effect | 3984 | 45 | 1433 | 1596 | 14.9 |
| Enhancing | 54 | 3 | 18 | 32 | 0.95 |
| **Total** | 4690 | 117 | 2108 | 2155 | 63.3 |

For each condition, we quantified the proteome mass fraction assigned to enriched GO biological processes. To avoid double-counting genes annotated to multiple processes each gene’s proteome mass was divided by the number of its GO annotations before estimating the contribution of each process. Figure 4 shows the GO processes with the largest proteome contribution within each fitness category under glucose and galactose growth conditions. In glucose, essential genes accounted for the largest fraction, representing 40% of the total proteome, whereas mean-effect genes contributed 14.9% (Table 1). Figure 4 shows the GO processes with the largest proteome contribution within each fitness category under both conditions. For readability, the visualization was restricted to the top five processes for essential, important, and mean-effect genes, and to the top two processes for the fitness-enhancing category.

**Figure 4.**
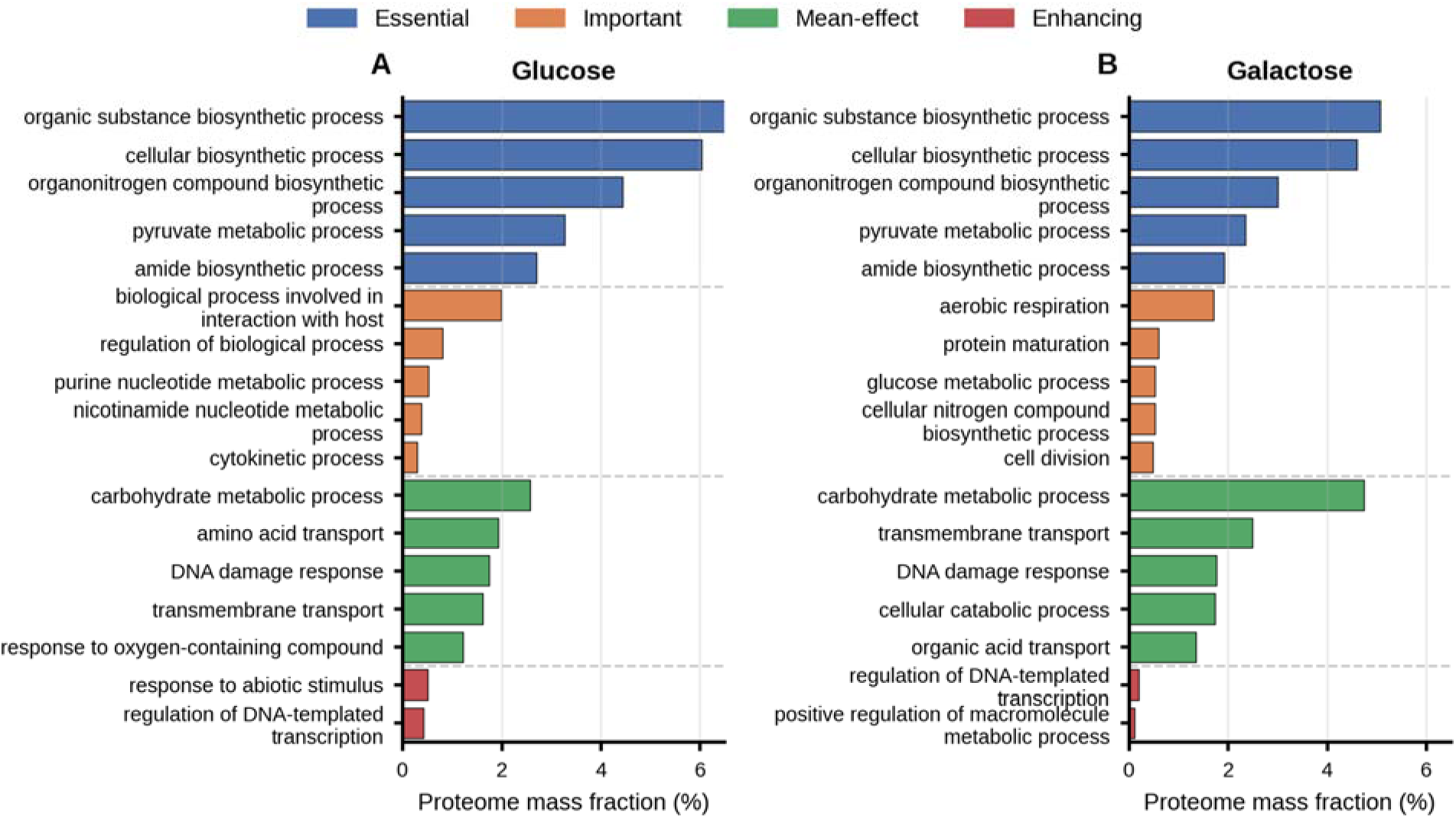
Enriched GO biological processes with the largest proteome mass fraction in glucose (A) and galactose (B) minimal media. Bars show proteome mass fraction, colors indicate fitness categories, and rows indicate GO terms. The top five GO processes are shown for Essential, Important, and Mean-effect categories, and the top two for Enhancing.

As shown in Fig. 4A-B, essential genes accounted for the largest proteome mass contribution in both glucose and galactose, mainly through biosynthetic processes. The important category contributed a smaller fraction and included more condition-specific functions, such as regulatory and nucleotide-related processes in glucose, and aerobic respiration, protein maturation, glucose metabolism, and cell division in galactose. Mean-effect genes were mainly associated with carbohydrate metabolism, transport, DNA damage response, and catabolic or stress-related processes, indicating that genes with little apparent fitness effect can still represent a substantial proteomic investment. The fitness-enhancing category showed the lowest contribution in both conditions, consistent with the limited number of genes in this group.

As mentioned above, our analysis revealed a subset of genes consistently classified as mean-effect across all conditions, thereafter referred to as core mean-effect genes. This subset shows a stable contribution to the proteome fraction around 0.11 of the total proteome fraction across conditions (Fig. 5A, Supplementary Table S14). An analysis of these 1946 core mean-effect genes provides further insights into potentially redundant, conditionally dispensable, or environmentally responsive functions. Several transport-related processes systematically appeared among the main GO terms ranked by proteome fraction, along with other stress-related functions, such as cellular response to DNA damage, which accounts for 0.47% of the total proteome (Fig. 5B).

**Figure 5.**
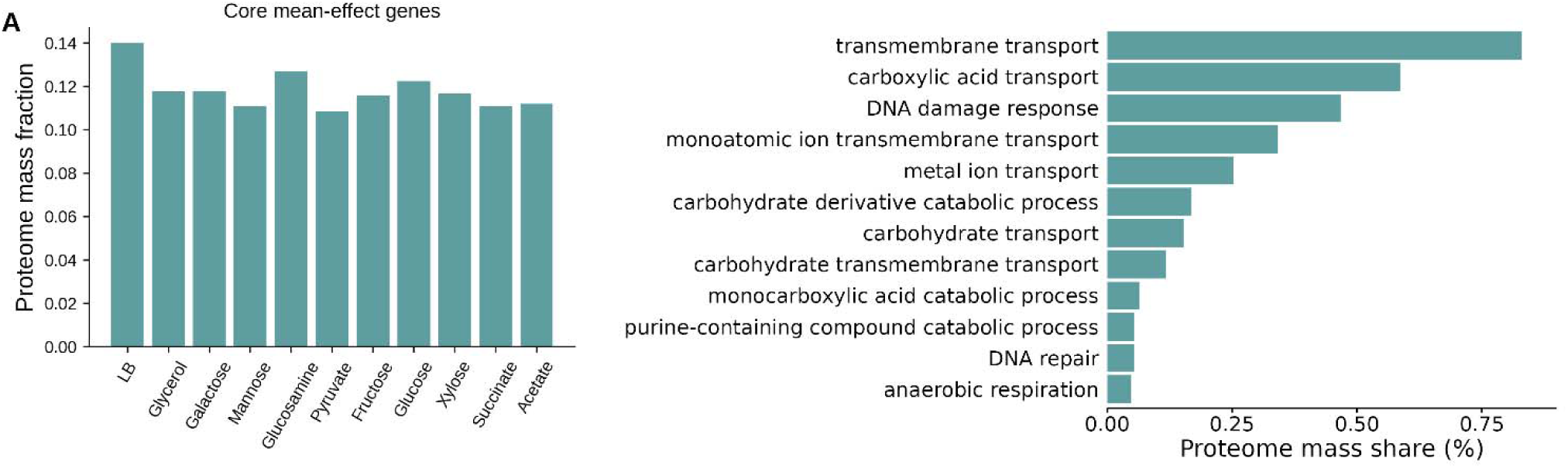
A) Core mean-effect proteome mass fraction under different conditions B) Proteome mass fraction from GO processes enriched in the core mean-effect genes under glucose minimal media condition.

### -Fitness enhancing genes are mostly transcriptional regulators

Genes belonging to the enhancing category refer to genes whose loss of function by deletion is beneficial to the cell as measured by a fitness increase (measured fitness value greater than 0.5). We initially considered whether this benefit might result from proteome savings, since often times a reduction in the synthesis of present but unused proteins could benefit fitness from a resource allocation perspective(O’Brien et al., 2016). However, as demonstrated earlier, the proteome fraction associated with the totality of fitness-enhancing genes remains negligible (Figure 4D). Across shared conditions, these genes were few and broadly overlapped with the genome-wide protein mass distribution (Supplementary Figure S2). To further explore the functions of the genes with the strongest fitness-enhancing profile, we identified those over a more stringent cut-off: a fitness value above 0.5 in at least 10 conditions, (n = 108). Performing a GO-term enrichment of this gene list for biological processes we found that twelve GO-terms were significantly enriched (Supplementary Table S15). The most significantly enriched GO term was ‘regulation of DNA-templated transcription’ (GO:0006355), followed by ‘response to abiotic stimulus’ (GO:0009628). Overall, the enriched biological processes primarily relate to transcriptional regulation, a finding supported by experimentally curated GO annotations (Gaudet et al., 2021)(15).

Some genes involved in sensing and regulation such as *rpoS, fis, hfq, arcB, greA, cspC, rcsB* are present in this list. Also, some small regulatory RNAs such as *oxyS, arcZ, gcvB* and *dsrA*. The gene encoding *rsd*, which regulates the activity of sigma D and makes the core RNAP more accessible for sigma S (Jishage et al., 2002), was shown to enhance fitness in non-stressful conditions. Other genes that were expected to enhance fitness in this category include, flagella (*fliT, flgN*) and ribosomal hibernation factors (*rmf, ettA*). Surprisingly, important functions such as ribosomal proteins (*rpsG, rpsR*), tRNAs (*selC, selU,valZ, thrT*), were also found in this category. These results are intriguing since the functions of these genes are in the essential gene categories. These genes would require further experimentation to confirm their fitness-enhancing properties and to elucidate the mechanism by which they enhance cellular fitness under the measured conditions.

Of special interest was the enrichment of transcriptional regulators within the fitness-enhancing category. Most bacterial transcription factors (TFs) help cells adjust to fluctuating environments. They bind signaling metabolites that alter their affinity for DNA. This, in turn, promotes or inhibits polymerase binding to specific genes, typically those tied to the use or production of the initial signal. We hypothesized that their fitness-enhancing properties could be explained by two possible mechanisms. First, under stable growth conditions, eliminating genes coding for regulators that respond to signals in conditions where such signals are absent should not be detrimental to the cell. This is because the effect of the TF over its regulated genes is not required under the condition tested. One such case is the *rpoS* gene, which encodes a sigma factor (sigma S or sigma 38) typically associated with stress responses, and plays a crucial role in regulating bet-hedging functions, survival mechanisms during stationary phase, and response to starvation. *rpoS* appears as a fitness-enhancing gene in seven out of ten growth conditions that do not involve stress. The second hypothesis is based on the observation that 54.1% of the genes in the *E. coli* Transcriptional Regulatory Network are regulated by two or more TFs, which allows for redundancy in their regulation. The loss of a redundant TF might imply the loss of fine-tuning in the modulation of a gene’s expression, but the remaining TFs are enough to block, or recruit polymerase, and the fluctuations in expression are not significant at the proteome level. An example of this is again *rpoS*, which regulates 340 genes, of which 209 are coregulated by other sigma factors (Fig. 6A)(Salgado et al., 2024). Eliminating the expression of all 340 regulated genes would free up proteomic resources equivalent to 15% of the total proteome. Whereas, eliminating only the 131 genes solely controlled by RpoS would free up 2% (Fig. 6B). This disparity, which is not proportional to the number of genes regulated, could underscore the relatively low contribution of the genes that are affected by a *rpoS* deletion in the conditions tested.

**Figure 6.**
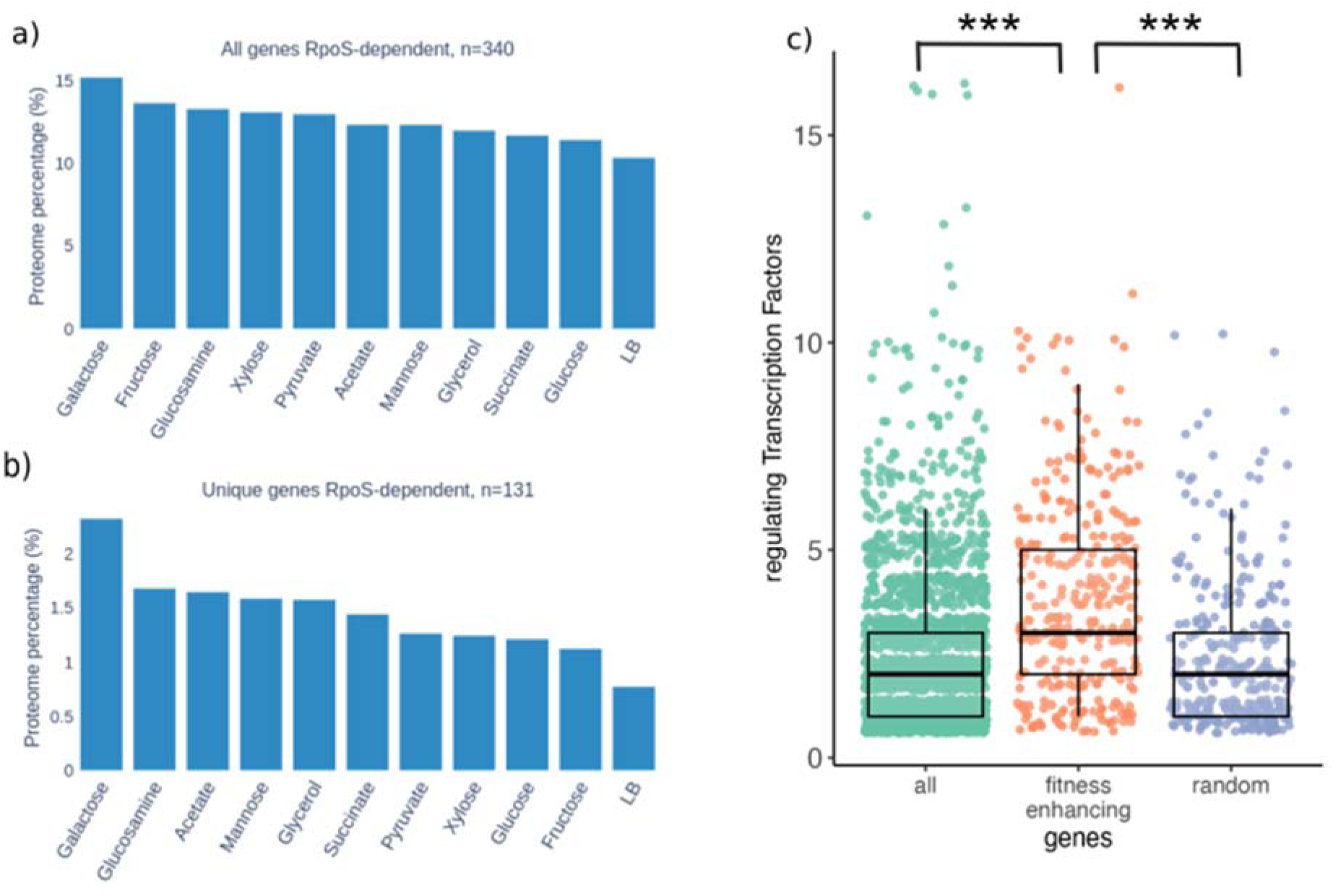
Fitness enhancing regulators. a) Calculation of the proteome percent that could be liberated by eliminating the expression of *rpoS* targets (sigma 38) in different carbon sources with shared targets, the genes identified for each regulon are listed in Supplementary Table S16. b) Potential proteome release distribution for the *rpoS* regulon with unique targets. c) The five fitness enhancing TFs regulate 323 genes that are on average coregulated by 3 other TFs this is significantly higher than the 2520 genes with known TF regulation and a random sample of genes (Wilcoxon-Mann-Whitney; p-value < 0.001).

The first hypothesis is difficult to validate systematically given our limited knowledge of the signals recognized by TFs (Ledezma-Tejeida et al., 2021). The second hypothesis is simpler to explore since redundancy in the regulation of a gene can be logically implied whenever a gene is regulated by 3 or more TFs, since there are only two possible effects (i.e. activation/repression), and the effect of the third regulator will necessarily be redundant to one of the other two. This regulatory redundancy requires a closer inspection in genes regulated by two TFs, since they could be having opposite effects over the transcription of the gene. Essentially, if our second hypothesis is correct, we would expect that fitness-enhancing TFs significantly regulate more genes also regulated by other TFs, and the deletion of a TF is not detrimental because others will still be contributing to the repression, or activation, of transcription. To test this, we looked for TFs that appear as fitness-enhancing in more than ten growth conditions, and five fit the cut-off criteria: RbsR, GcvA, Fis, SlyA, and RcsB. Together, they regulate 323 genes, which are coregulated by 3 other TFs, on average. To test whether this level of coregulation is expected within the TRN, or is significantly higher, we obtained the number of TFs that regulate each of the 2521 genes present in the network (Salgado et al., 2024), and a random sample of 323 genes to rule-out effects from the sample size. The full network, and the random sample of genes showed a similar percentage of genes regulated by 3 or more TFs: 31.4% and 35.2% respectively, are genes with implied regulatory redundancy. In contrast, 68.4% of genes regulated by fitness-enhancing TFs show implied redundancy (Table 2, Supplementary Figure S6). To further support this result, and include genes regulated by 2 TFs in the analysis, we evaluated whether any pair of TFs regulating the same gene had the same effect over its expression. In total, 43.2% of the genes in the TRN, and 45.2% of the genes in the random sample showed regulatory redundancy, whereas among the genes regulated by fitness-enhancing TFs the percentage rose to 72.7% (Table 2). Finally, we included the genes regulated by one TF in each dataset, and compared the distributions of the number of TFs regulating each gene in the three datasets (Figure 6C). Results show that the genes regulated by fitness-enhancing TFs are significantly regulated by more TFs (Wilcoxon-Mann-Whitney; p-value < 0.001). These results support our hypothesis that regulatory redundancy contributes to the fitness-enhancing quality of these regulators.

**Table 2.** Gene counts within the complete Transcriptional Regulatory Network (TRN), a random sample of 323 genes to account for sample size biases, and the 323 genes regulated by the Fitness/enhancing Transcription Factors (TFs).

|  | TRN | Random sample | Fitness-enhancing |
| --- | --- | --- | --- |
| **Genes regulated by 1 TF** | 1157 (45.9%) | 139 (43%) | 58 (17.9%) |
| **Genes regulated by 2 TFs** | 571 (22.6%) | 70 (21.6%) | 44 (18.9%) |
| **Genes regulated by 3 or more TFs** | 792 (31.4%) | 114 (35.2%) | 221 (68.4%) |
| **Total** | 2520 | 323 | 323 |
| **Genes with regulatory redundancy** | 1089 (43.2%) | 146 (45.2%) | 235 (72.7 %) |

### Validation of fitness benefits with a long-term experimental evolution experiment

In an experimental evolution setting, bacterial populations who survive by adapting consistently exhibit long-term increases in fitness. Such evolutionary frameworks provide a rigorous test of the generalizability of insights obtained from systematic single-gene knockout and protein expression analyses. To compare and cross-validate our findings on fitness and classifications, we utilized data from long-term experimental evolution (LTEE) in *Escherichia coli* (Tenaillon et al., 2016). The LTEE offers a unique perspective for observing sustained adaptive changes over tens of thousands of generations.

The ALEdb (Phaneuf et al., 2019) compiles and classifies the mutations present in populations of the LTEE experiment after 50,000 generations of experimental evolution in a minimal glucose environment (Tenaillon et al., 2016) (LTEE ARA). First, we examined the spectrum of genomic mutations, regardless the type, it revealed that the majority were associated with mean fitness consequences in a non-evolved strain (Supplementary Figure 4A). This analysis revealed that most mutations are around zero values in measured fitness (Supplementary Figure 4A). We then filtered our datasets for putative loss-of-function mutations such as frameshift, nonsense, large deletion, and insertion sequence disruptions. From 459 annotated genes with loss-of-function mutations, 423 are mean-effect, 13 are essential 12 are important and 11 are fitness enhancing (Supplementary Figure 4B). Transporter genes comprised a prominent subset of loss-of-function mutations: 62 transporters were disrupted in the LTEE_ARA dataset. Consistent with the fitness assays, nearly all (57/62) displayed mean fitness values in glucose minimal medium, while only five showed deleterious effects (Supplementary Figure 4C). Among transcriptional regulators, we observed parallel loss-of-function mutations, most notably in *rpoS* and *malT*. Of the 29 regulator genes harboring loss-of-function mutations, 28 are classified as mean or beneficial in its fitness effects (Supplementary Figure 4D), with the largest benefit observed for *rpoS* (+1.00) and *yebK* (+0.57); only *allR* displayed a notable detrimental effect (−0.541).

Regarding gene expression changes observed in the LTEE, we find patterns that are consistent with our results. Favate et al., 2022 (Favate et al., 2022) showed that parallel mutations in global regulators such as *rpoS*, *hns*, and *nadR* rewire transcriptional programs, reducing costly bet-hedging functions, stress responses, and unused transport systems. In line with this, Mori et al., 2024 (Mori et al., 2024) demonstrated that these regulatory changes manifest at the proteome level as a smaller fraction of proteins that remain inactive or unnecessary under the growth conditions. This includes the down-regulation of amino acid biosynthetic enzymes (ArgA, IlvA/E), molecular chaperones (GroEL/GroES), and dispensable transporters, highlighting that fitness gains arise not from improved enzyme kinetics but from more efficient proteome allocation. Overall, these results indicate that selection in the LTEE favors genotypes that improve fitness by reducing the production of costly proteins that provide no benefit in the experimental conditions.

### Many fitness-enhancing genes have unknown functions

We identified a subset of genes that fall into the fitness-enhancing category in the majority of the 46 fitness conditions (fitness conditions not necessarily shared with the proteomics dataset). When we looked at the top 100 genes of this subset we found that 28% had unknown functions. In an effort to define the function of these genes, we leveraged Artificial Intelligence (AI) and a hidden Markov model profile comparison to present a homology-based functional hypothesis for these genes. Our annotation process heavily relied on homology, where similarities in sequence or structure suggest a shared ancestry between proteins, enabling a testable hypothesis of transfer of function from experimentally determined proteins to targets (Fig. 7).

**Figure 7.**
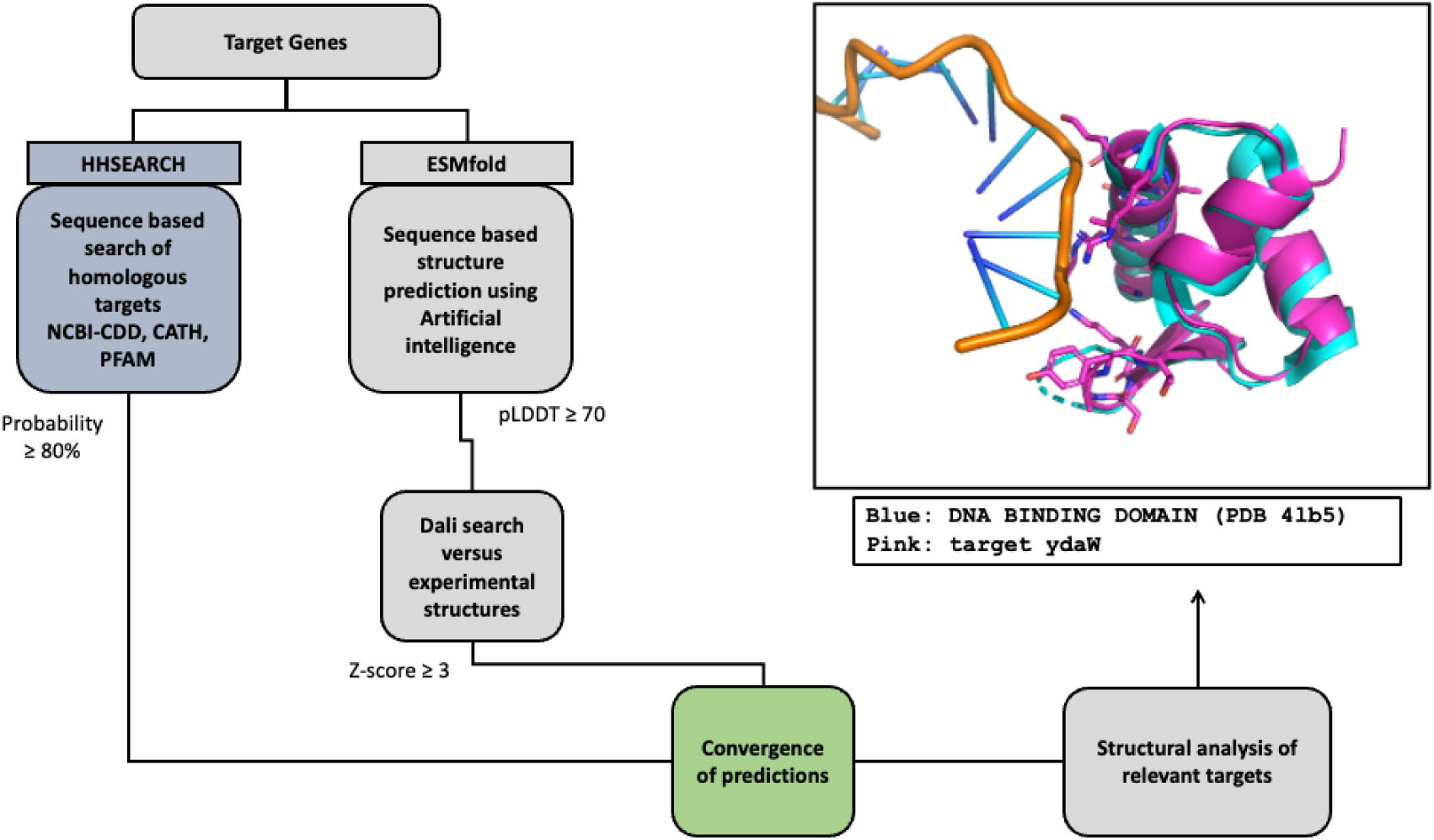
An AI assisted bioinformatic workflow to perform a functional inference of 28 fitness-enhancing genes by a combination of sequence and structure-based approaches. As an example, the b1361 target was annotated as a DNA binding protein based on high sequence and structure similarities with two DNA binding proteins (PDB ids: 7bzh and 4lb5). The figure displays a structural alignment between the structure of the DNA dependent protein kinase PKZ in complex with DNA (PDB 4lb5) in gray and the structural model of b1361 produced with AlphaFold2 (in pink).

For the structure and sequence-based functional inference we applied a combination of sequence and structure-based approaches. We built Hidden Markov Models (HMM) for all targets using the HHsearch suite implemented in the MPI toolkit (Gabler et al., 2020) and we employed them to search the NCBI conserved domains (Wang et al., 2023), PFAM (Blum et al., 2025) and PDB databases (Berman, 2000), represented by HMM profiles. In parallel we built structural models for the targets using the AI algorithms for structural prediction ESMFold (Lin et al., 2023) and AlphaFold2 (Mirdita et al., 2022). We compared the structural models with the whole PDB database employing the DALI server (Holm et al., 2023). As an example of the annotation process we describe how we annotated the sequence of the b1361 target (Fig. 7). We first created an HMM for the target and searched the PFAM, CDD and PDB databases. The query protein hit the sequence of the 7bzh structure with high probability (98.29) the length of the alignment was 51 residues with an E-Value: 1.3e-5. This protein was annotated as a DNA-binding protein in the PDB database. Next, with the same sequence we produced an AI-structural model that was superimposed with the crystal structure of the DNA dependent protein kinase PKZ in complex with DNA (PDB 4lb5). The structural alignment displayed scores that imply structural homology: Z-score=9.0, rmsd 1.0 Å for 52 alpha carbons. Since the target structure was determined in complex with DNA, these results support the hypothesis that b1361 may function as a DNA binding protein. From the 28 proteins with unknown function, we present predictions of 15 targets, with good confidence based on high scores for both the sequence-based approach (probability ≥ 90%) and the structure-based approach (Z-score ≥ 8). We also present predictions of functions of 9 targets, with lower confidence (lower scores for the predictions, with details in Supplementary Table S17). And we could not make predictions for 4 targets. We found four putative transcriptional regulators, four putative metabolite-binding proteins, three putative DNA-binding proteins and three putative lipoproteins among the targets. These predictions require experimental confirmation (Supplementary Table S17). These results show that most of our functional hypotheses for previously uncharacterized fitness-enhancing genes are related to the regulation of gene expression. Among the 15 high confidence predicted targets, almost half of them may have a regulatory role. Four of them are predicted transcriptional regulators and the other three proteins have a predicted DNA binding motif that may indicate regulatory function.

### Comparing *E. coli* fitness data to other organisms

In order to generalize our findings to other organisms we analyzed data from *Pseudomonas putida* KT2440 where fitness data(Price et al., 2018) and expression profiles are available (Bojanovič et al., 2017) (28) in the same growth condition. We classified *P. putida* genes into the same fitness categories used for *E. coli* (essential, important, mean-effect, and fitness-enhancing) applying identical fitness cutoffs. From the expression profiles, we identified the top 5% of expressed genes, which corresponded to the 100 most highly expressed genes. Our analysis focused on the highly expressed genes within the mean-effect and fitness-enhancing categories. This subset included eight transcriptional regulators, various stress-related genes (e.g., those involved in copper resistance, cold shock response, and chaperone functions), and several genes with unknown functions (Supplementary Figure S5). Interestingly, unlike *E. coli*, this list in *P. putida* also included several metabolic genes. This highlight both shared mechanisms, such as stress response, and species-specific functions, such as metabolic functions in *P. putida*, reflecting its renowned metabolic flexibility (Belda et al., 2016).

We further analyzed fitness profiles across nine bacterial species: three alphaproteobacteria (*Caulobacter crescentus, Dinoroseobacter shibae, Sinorhizobium meliloti*), five gammaproteobacteria (*Pseudomonas fluorescens, Pseudomonas simiae, Pseudomonas putida*, *Pseudomonas stutzeri, Escherichia coli*), and one representative of the Bacteroidetes (*Echinicola vietnamensis*) (Price et al., 2018) (Supplementary Figure S6). Across the nine species, GO-based functional categorization revealed similar patterns among mean-effect and fitness-enhancing genes. Motility and transport functions were frequently represented within these groups. Our multi-species comparative analysis (Supplementary Figure S5) suggests that motility, transport, and stress response may represent fitness-relevant adjustments under nutrient limitation. However, an important caveat is that, for many species in this study, the absence of matched gene expression data precludes quantification of the expression costs associated with these potential hedging traits.

## Discussion

A generalist bacterium has a broad niche: it can reproduce under changing environments, utilize diverse substrates, and withstand multiple abiotic stress challenges (Zhu and Dai, 2024)(Albalat and Cañestro, 2016). The adaptive regulation of gene expression paradigm assumes that conditions are static, however this is rarely the case in natural environments (Mitchell et al., 2009). Random Barcoded transposon sequencing (RB-Tnseq) experiments provide unprecedented quantitative fitness data of mutants in many environments. In this work, we used publicly available datasets consisting of fitness and proteomic scores of *Escherichia coli* grown across several static conditions to study the quantitative relationship among expression costs and fitness contribution of a gene.

We found that a large fraction of genes presents a mean fitness effect in the static environments we analyzed; genes with mean fitness effects account for up to 75% of the proteome in rich medium (LB) and up to 60% in minimal medium (glycerol; Fig. 1C). Similar magnitudes have been reported in other bacteria. For example, in the marine bacterium *Ruegeria pomeroyi* DSS-3, approximately 50% of the theoretical proteome is expressed only under specific environmental conditions (Christie-Oleza et al., 2012), highlighting that the production of condition-specific proteins is common across diverse microbes. To determine which functions dominate this “unused” fraction, we performed gene-function enrichment analyses for each fitness category and found that transport functions are strongly overrepresented. These transport systems are often expressed because they prepare the cell for potential future changes in nutrient availability, even if they provide no immediate benefit in the current environment (anticipatory mechanism). This interpretation aligns with previous work showing that transporters exhibit one of the largest discrepancies between model-predicted proteome allocation and experimentally measured proteome levels (Hu et al., 2023). We corroborate this trend from a different angle: transporters comprise one of the largest groups of genes with no mean fitness effect in our analyses. Based on these results, we hypothesize that reducing the expression of transport systems that are unnecessary in a given environment frees proteomic resources, as well as membrane space, which can instead support functions essential for growth, such as respiratory chain components.

Our analysis of fitness enhancing genes showed that regulatory functions, including transcriptional regulation, were among the most prevalent functional categories. Furthermore, among enhancing genes of unknown function, structural analyses suggested that 46% of the highest-scoring candidates may encode putative transcriptional regulators. Remarkably, *rpoS*, the global stress related sigma factor has experimental evidence to be fitness enhancing, consistent with other studies (Mata et al., 2017; Bouillet et al., 2024). One study showed that *rpoS* mutant strains have a competitive growth advantage, possibly due to a greater nutrient uptake capacity (32). In another study, a strain engineered to produce putrescine increased product yield in a deletion mutant (Qian et al., 2009). Surprisingly, transcriptomic analysis showed that RpoS also represses important genes, such as those related to the flagellum and several enzymes in the TCA cycle (Patten et al., 2004). Modulating *rpoS* expression holds significant potential for generating strains with novel and interesting phenotypes, since it is not considered essential. However, *rpoS* mutants have been demonstrated to have a poor survival capacity in stressful conditions (Schellhorn, 2020).

Our results show there is a cost associated with the expression of stress anticipatory mechanisms (see above). In the case of *rpoS,* this cost can be quantified in terms of proteome (up to 15% of the proteome in galactose; Figure 6A). This anticipatory mechanism is encoded in the transcriptional regulatory network, and it has a quantifiable fitness cost. It has been previously shown that beneficial null mutations mostly target regulatory functions (Hottes et al., 2013). In a previous study, our group showed that eliminating transcriptional regulators that activate the expression of unused functions increased the proteomic budget of an engineered *E. coli* strain(Lastiri-Pancardo et al., 2020) Here, we show that much of the proteome can be reallocated by engineering regulatory programs that reduce the expression of anticipatory functions. We found that fitness-enhancing transcription factors (TFs) regulate genes with significantly more co-regulators than other genes in the network. This suggests that their fitness-enhancing effect is unlikely to stem from proteome savings due to the elimination of the TF itself, but rather from suboptimal regulation in static environments, where redundant or overlapping TFs may compensate for the loss of fine-tuned control.

Together, our LTEE analysis and multi-species comparisons support this conclusion. In the LTEE, our results are consistent with fitness gains arising from resource reallocation rather than improved enzymatic catalysis, as reflected by the fixation of loss-of-function mutations. This theme extends beyond a single species, as our comparative analysis suggests that processes such as motility and transport are recurrent, fitness-relevant adjustments to resource investment under resource constraints.

This analysis has several limitations, derived from the nature of the datasets used, and presents several opportunities for further exploration. First, experimental measurements are noisy and inherently contain errors. Second, because neither dataset covers all genes, using only their overlap excludes a substantial fraction of genes from the analysis. Third, the data in this study are derived exclusively from mid-log phase growth conditions, predominantly in minimal medium with a single carbon source. Although these conditions are standard in laboratory settings, they do not accurately represent the natural environments in which bacteria have evolved. Furthermore, using single-gene deletion data does not provide information about genetic interactions. Such unintuitive and often unpredictable interactions could potentially arise through the elimination of the functions of one or more genes. This is particularly relevant for genes that encode for proteins with the same function, such as isoenzymes.

## Closing remarks

We identified genes that increased fitness when deleted and discovered that, in a wild type strain, they have low levels of expression. Many of these genes had unknown functions, and were often regulated by *rpoS*, a general stress sigma factor that controls up to 15% of the proteome. To elucidate the main functions of the genes in each category, we assigned them functional annotations. Finally, we calculated the proteome mass fraction and the proteome elimination potential of each category and compared them with the results from the ME-model simulations. We found that up to 14% of the proteome consisted of genes whose disruption did not severely affect fitness. This zero-base budget exercise provides estimates of the expression costs associated with genes and cellular functions with non-severe fitness effects across different environments.

## Statements

### Data availability statement

Github repository with the code and source data to reproduce this analysis: https://github.com/utrillalab/Fitness_analysis

## Acknowledgements

We acknowledge Dre. Clau Hernandez Armenta and the UNAM Postdoctoral Program (POSDOC) 2021_2 for initial contributions to this project supervised by JU. We acknowledge technical support by Dra. Gabriela Perez-Segura

## Author contributions

D.H. and J.U. designed research. D.H., L.S-A., O.A.A-V., D.L-T., J.F-R. and J.U. performed research. D.H., L.S-A., O.A.A-V., and J.F-R. analyzed data. D.H. D.L-T and J.U. performed critical data analysis. J.U. supervised and guided the research. D.H., LS-A. and J.U. wrote the manuscript.

