## supplemental material for "Potential benefit of loss-of-function on bacterial fitness"

**Supplementary Material**

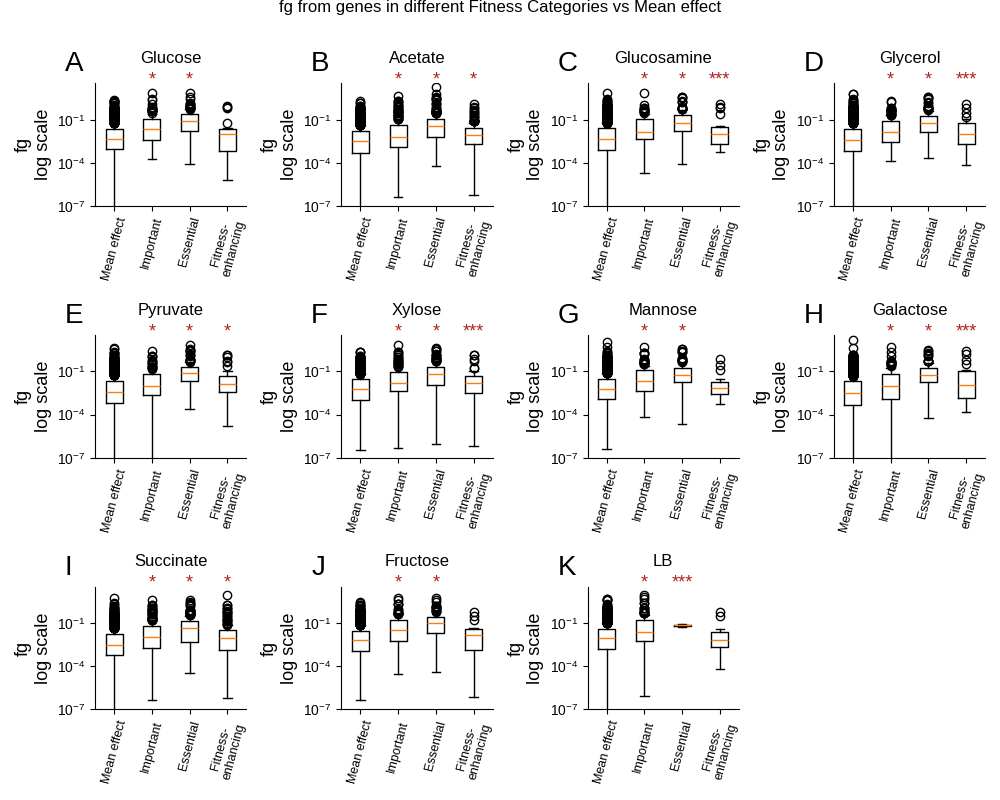

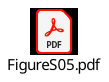

Figure S1 (Related to Fig. 1). Distribution analysis for each fitness category’s protein mass contributions. A-K, Box plots of each Fitness category’s protein mass contributions in all shared conditions. Statistical significance based on a Kruskal-Wallis test is denoted by: * (99.9% confidence) and *** (90% confidence).

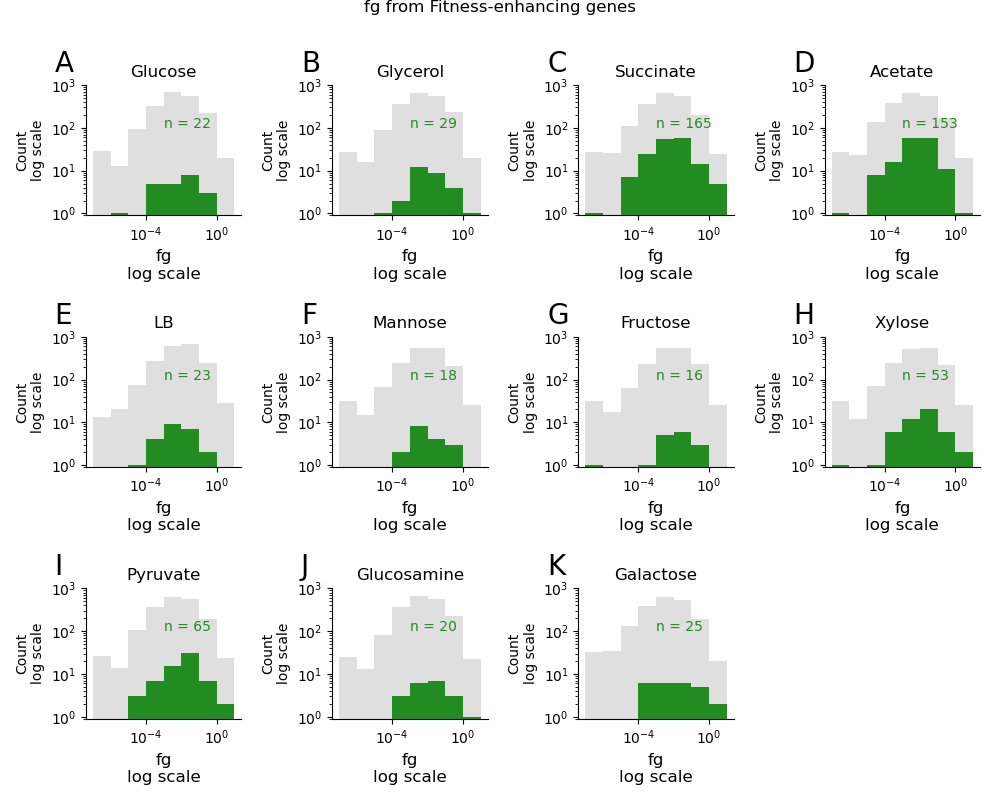

Figure S2 (Related to Fig. 1). Distribution of the Fitness-enhancing genes in the eleven shared conditions of proteomics and fitness. Grey shadows show the distributions of all genes. Green distributions correspond to those of fitness-enhancing genes

Figure S3 (Related to Figure 6). Distribution of Transcription Factors regulating genes in (a) the full Transcriptional Regulatory Network, (b) a set of 323 random genes taken from the network, and (c) the 323 genes regulated by fitness-enhancing Transcription Factors. Y axes are shown as percentages for easier comparison among panels.
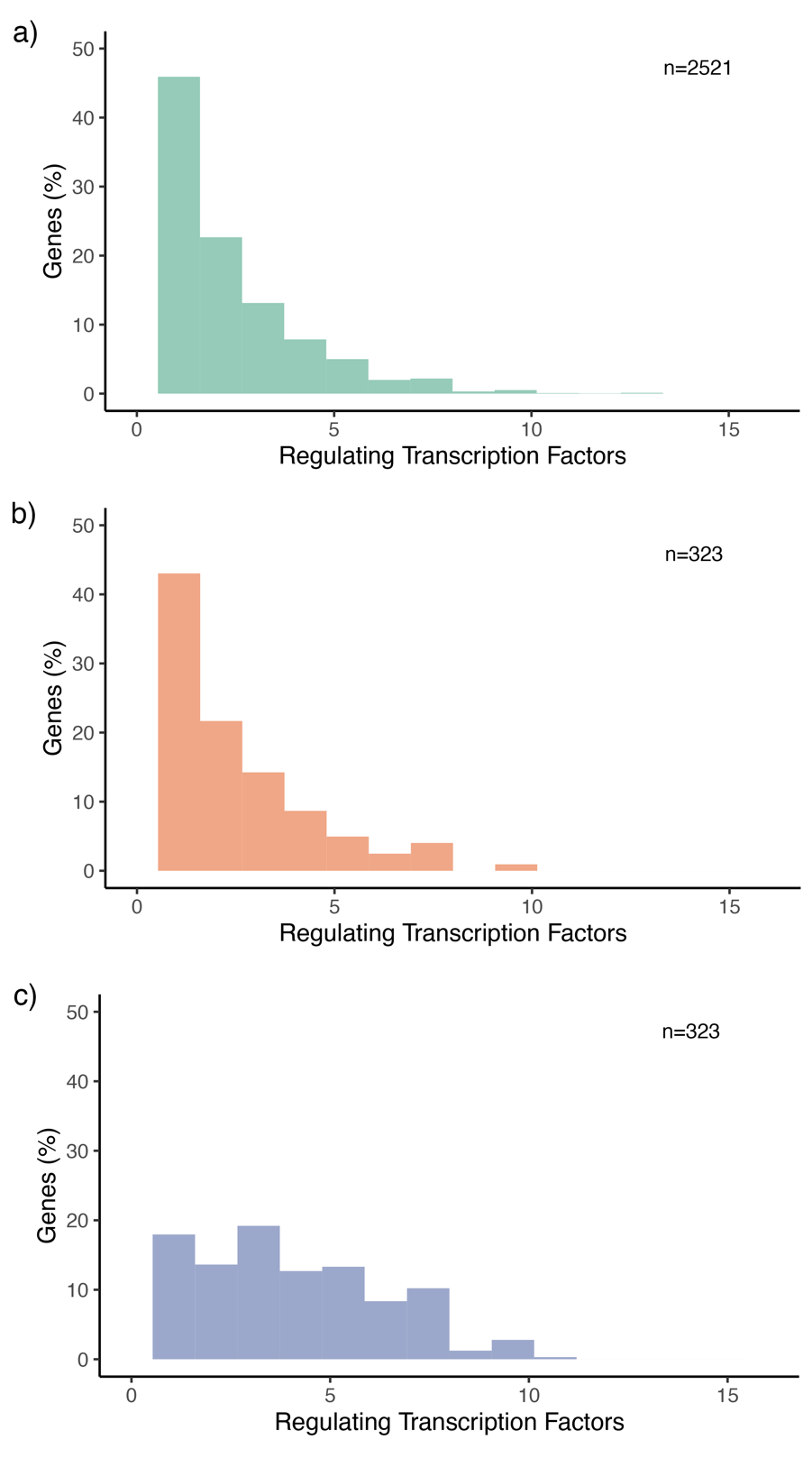

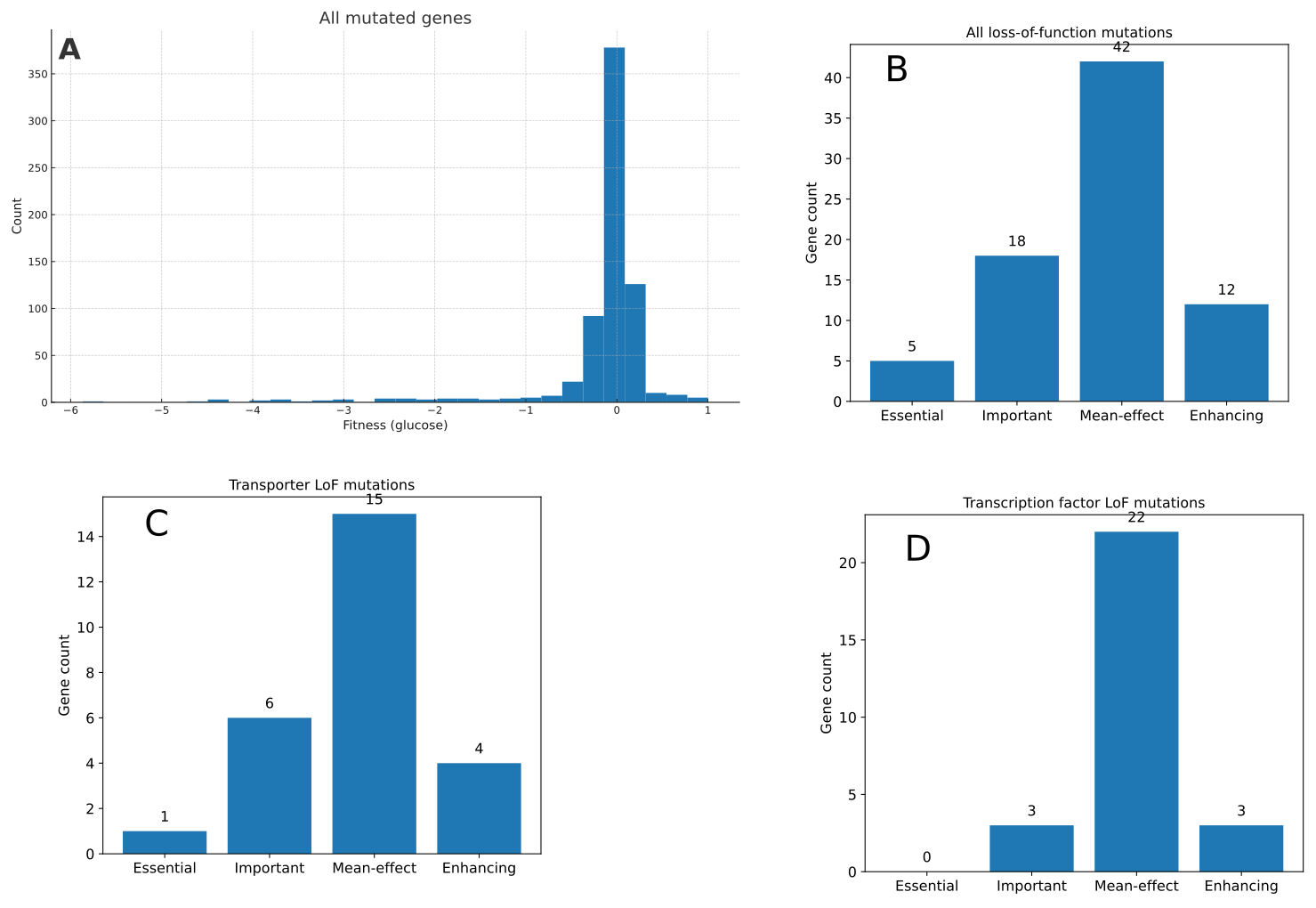

Figure S4 Analysis of LTEE_Ara mutations A) fitness values of all mutated genes B )Fitness classifications of Loss-of-function mutations C) Fitness classifications of genes annotated as transporters in all loss-of-function mutations (LoF) D)Fitness classifications of genes annotated as Transcription factors in loss-of-function (LoF) mutations

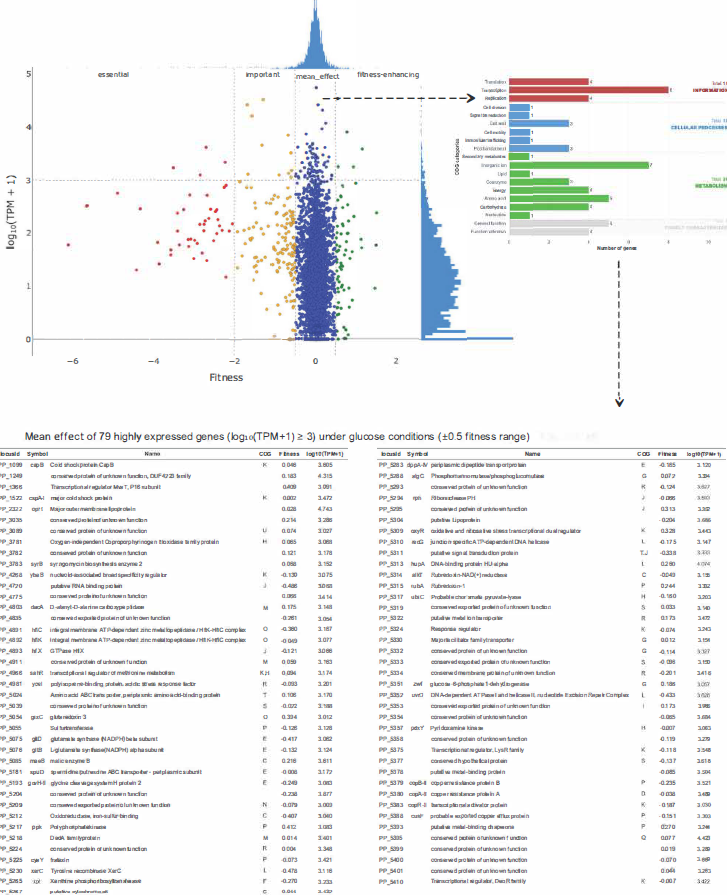

Figure S5. Fitness versus expression in *Pseudomonas putida,* the 79 most expressed genes that are in the no effect category are showed in the table.

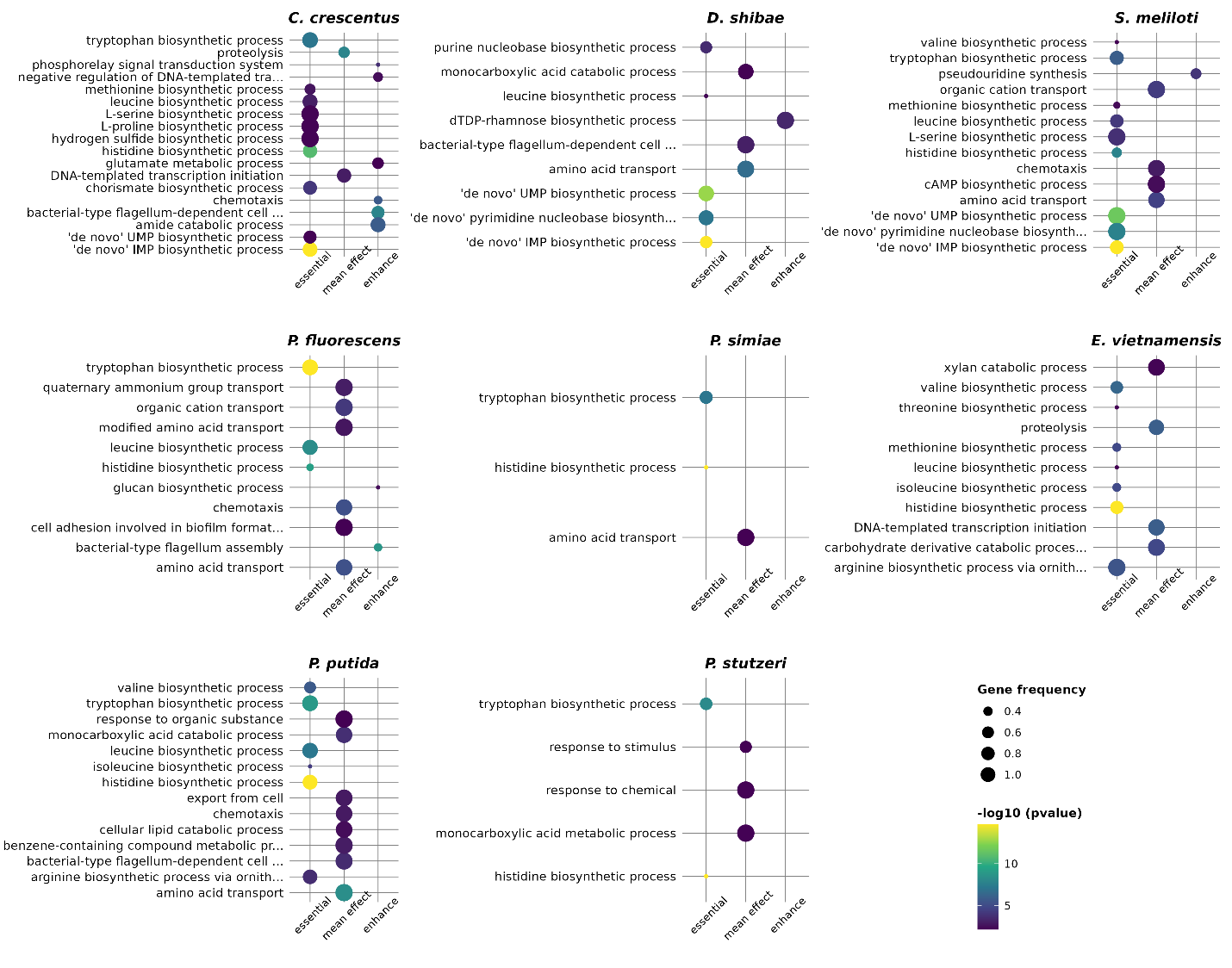

Sfigure 6. GO term enrichment analysis of genome-wide mutant fitness profiles across nine bacterial species: three alphaproteobacteria (*Caulobacter crescentus*, *Dinoroseobacter shibae*, *Sinorhizobium meliloti*), five gammaproteobacteria (*Pseudomonas fluorescens*, *Pseudomonas simiae*, *Pseudomonas putida*, *Pseudomonas stutzeri*, *Escherichia coli*), and one representative of the Bacteroidetes (*Echinicola vietnamensis*). Enriched GO categories are shown for each gene fitness class, except for the *Important* category, which was excluded due to its variability and to simplify interpretation.

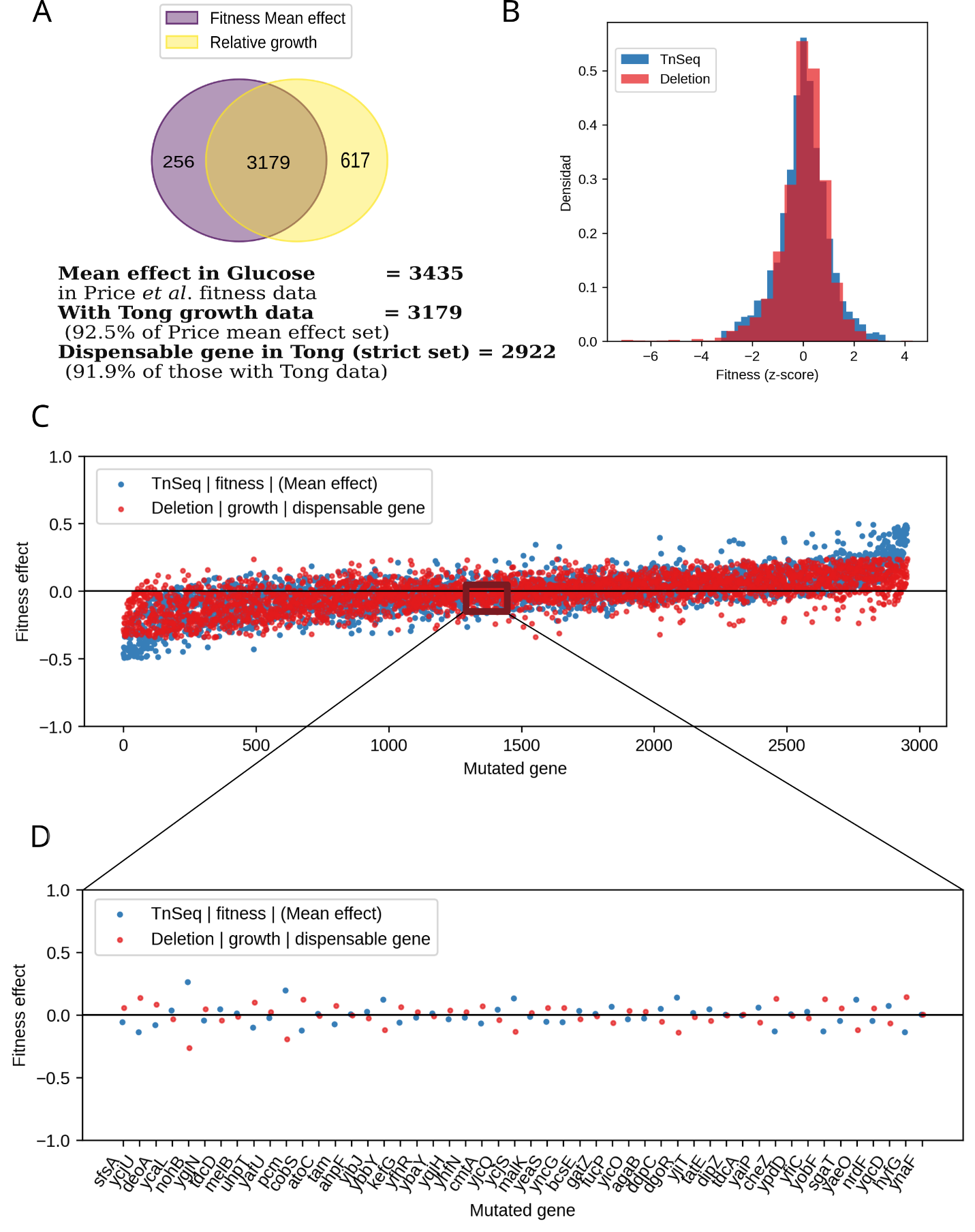

**Figure S7. Consistency between pooled TnSeq fitness and single-deletion growth for genes classified as “no-effect” in glucose.**

**(A)** Venn diagram showing overlap between genes with no-effect fitness in glucose in the Price pooled TnSeq dataset and genes with relative growth measurements in the Tong single-deletion dataset. Numbers indicate genes unique to each set and shared between them. Text below reports the total number of no-effect genes in glucose in Price, the subset with Tong growth data, and the number that are also classified as neutral in Tong under a strict neutrality threshold.

**(B)** Overlaid histograms of standardized fitness (z-scores) for the shared no-effect genes, comparing TnSeq fitness (blue) and deletion growth (red).

**(C)** Fitness effect for each shared no-effect gene, ordered along the x-axis by gene index. Blue points correspond to TnSeq fitness values and red points to deletion growth values; the horizontal line indicates zero effect. The boxed region marks a subset of genes highlighted in panel D.

**(D)** Zoom-in of the boxed region from panel C, showing individual gene symbols on the x-axis and the corresponding TnSeq (blue) and deletion (red) fitness effects.

**Supplementary table 17. Functional annotation of gene targets sorted by size**

The analysis of 28 genes confidently predicted the function of 15 targets with high HHsearch probability (>90%) and Z-scores exceeding 8. Nine targets were predicted with low confidence, while four had no homologous hits and/or were structurally predicted with low confidence. Only structural models with better than 70 predicted local distance difference tests (pLDDT) were used in the analysis. Among the predicted functions were enzymes, proteases, ion transporters, lipid-binding proteins, endonucleases, DNA-binding proteins, transcriptional regulators, and translation repressors. Notably, targets of viral origin were also identified.

| **Confidence**  **Level** | **Protein target**  **Annotated in EcoCyc** | **Annotated by structure** | **Annotated by sequence** | **Functional Annotation**  **Structural model** |
| --- | --- | --- | --- | --- |
| **High confidence**  **Annotation** | b0955 sp\|P75867\|LONH_ECOLI  Structure 07  gene: ycbZ  polypeptide: putative ATP-dependent protease YcbZ | Hit with: 3k1j  z-score: 25.1  rmsd:3.2  Aligned residues:497 | Hit with: 3k1j  Probability: 99.97  Aligned columns: 478  E-Value: 9.7e-28 | Several domains:  ATP-dependent protease Lon; ATP-binding; Nucleotide binding  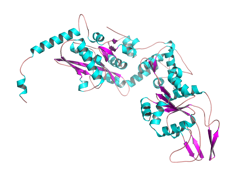 |
| **High confidence**  **Annotation** | b1754 sp\|P76223\|YNJB_ECOLI  Structure 023  gene: ynjB  polypeptide: putative ABC transporter periplasmic binding protein YnjB | Hit with: 6dtq  z-score: 27.7  rmsd:2.9  Aligned residues:326 | Hit with: 8c8f  Probability: 100  Aligned columns: 355  E-Value: 2.1e-33 | Maltose/maltodextrin-binding periplasmic protein  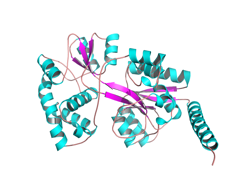 |
| **High confidence**  **Annotation** | b1422  Structure 028  sp\|P77171\|YDCI_ECOLI  gene ydcI  polypeptide: DNA-binding transcriptional dual regulator YdcI | Hit with: 3fxq  z-score: 20.1  rmsd:3.2  Aligned residues:227 | Hit with: 3fxq  Probability: 100  Aligned columns: 296  E-Value: 3.5e-52 | LysR-type transcriptional regulator TsaR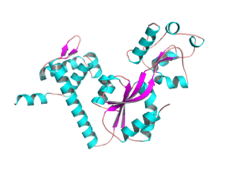 |
| **High confidence**  **Annotation** | b0658 sp\|P0AE78\|CORC_ECOLI  Structure 018  gene:ybeX  polypeptide: CorC-HlyC family protein YbeX | Hit with: 3nqr  z-score: 22.7  rmsd:1.3  Aligned residues:123 | Hit with: 4hg0  Probability: 100  Aligned columns: 291  E-Value: 7.6e-58 | magnesium and cobalt efflux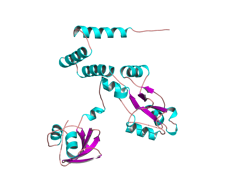 |
| **Low confidence**  **Annotation** | b3644 sp\|P23839\|YICC_ECOLI  Structure 013  gene: yicC  polypeptide: putative RNase adaptor protein YicC | Hit with: 8ax9  z-score: 5.4  rmsd:2.2  Aligned residues:73 | Hit with: 8ax9  Probability: 90.59  Aligned columns: 154  E-Value: 37 | Lipid binding protein  N-terminal domain  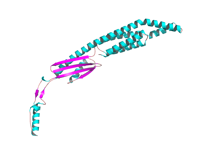 |
| **High confidence**  **Annotation** | b1725 sp\|P77739\|KT3K_ECOLI  Structure 012  gene: yniA  polypeptide: putative kinase YniA | Hit with: 3jr1  z-score:38.4  rmsd:1.5  Aligned residues:285 | Hit with: 3jr1  Probability: 100  Aligned columns: 285  E-Value: 9.5e-30 | Fructosamine kinase  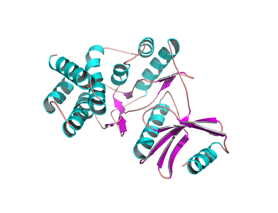 |
| **High confidence**  **Annotation** | b2229 sp\|P76466\|YFAT_ECOLI  Structure 02  gene:yfaT  polypeptide: DUF1175 domain-containing protein YfaT | Hit with: 3g27  z-score:16.7  rmsd:0.7  Aligned residues:81 | Hit with: 7Bk8  Probability: 99.85  Aligned columns: 186  E-Value: 3.7e-19 | bacteriophage protein  Peptidoglycan binding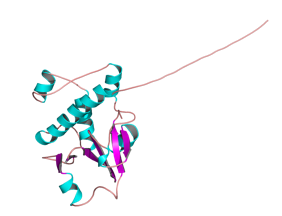 |
| **High confidence**  **Annotation** | b1758 sp\|P76226\|YNJF_ECOLI  Structure 026  gene:ynjF  polypeptide:CDP-alcohol phosphatidyltransferase domain-containing protein YnjF | Hit with: 5d92  z-score:20.6  rmsd:2.1  Aligned residues:188 | Hit with: 6h59  Probability: 99.89  Aligned columns: 196  E-Value: 4.4e-20 | Phosphatidylinositol Synthase  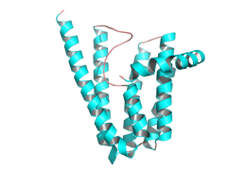 |
| **High confidence**  **Annotation** | b0190 sp\|P0AA97\|YAEQ_ECOLI  Structure 024  gene:yaeQ  polypeptide: uncharacterized protein YaeQ | Hit with: 3c0u  z-score:30.8  rmsd:1.6  Aligned residues:177 | Hit with: 3C0U  Probability: 100  Aligned columns: 181  E-Value: 2.5e-54 | Endonuclease  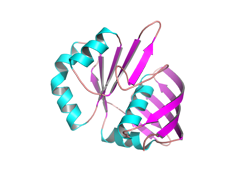 |
| **High confidence**  **Annotation** | b2948 sp\|P0A8W5\|YQGE_ECOLI  Structure 015  gene:yqgE  polypeptide:DUF179 domain-containing protein YqgE | Hit with: 2haf  z-score:26.8  rmsd:1.4  Aligned residues:185 | Hit with: 2haf  Probability: 100  Aligned columns: 183  E-Value: 5.1e-4 | Translation repressor  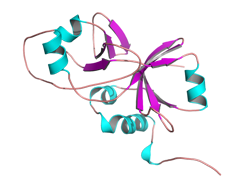 |
| **High confidence**  **Annotation** | b1361 sp\|P76066\|YDAW_ECOLI  Structure 019  gene:ydaW  polypeptide: Rac prophage; putative uncharacterized protein YdaW | Hit with: 4lb5  z-score:9.0  rmsd:1.0  Aligned residues:52  **DNA bound in target** | Hit with: 7bzh  Probability: 98.29  Aligned columns: 51  E-Value: 1.3e-5 | DNA Binding Protein  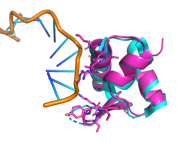 |
| **High confidence**  **Annotation** | b2123 sp\|P33354\|YEHR_ECOLI  Structure 022  gene:yehR  polypeptide:DUF1307 domain-containing lipoprotein YehR | Hit with: 2joe  z-score:16.5  rmsd:1.8  Aligned residues:126 | Hit with: 2joe  Probability: 99.94  Aligned columns: 131  E-Value: 6.6e-24 | Lipoprotein  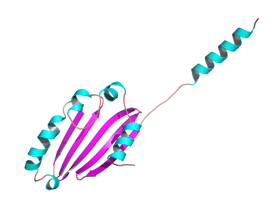 |
| **High confidence**  **Annotation** | b2446 sp\|P76546\|YFFO_ECOLI  Structure 08  gene: yffO  polypeptide: CPZ-55 prophage; uncharacterized protein YffO | Hit with: 4dyq  z-score:11.3  rmsd:3.4  Aligned residues:100 | Hit with: 4dyq  Probability: 99.54  Aligned columns: 111  E-Value: 1.7e-12 | Terminase Subunit  DNA-binding  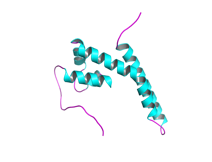 |
| **High confidence**  **Annotation** | b3611 sp\|P0AG27\|YIBN_ECOLI  Structure 016  gene:yibN  polypeptide:putative sulfurtransferase YibN | Hit with: 1gmx  z-score:17.1  rmsd:2.3  Aligned residues:107 | Hit with: 1yt8  Probability: 99.38  Aligned columns: 103  E-Value: 2.8e-11 | Thiosulfate sulfurtransferase  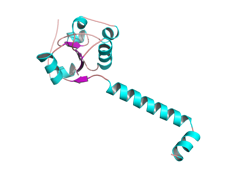 |
| **Low confidence**  **Annotation** | b0468 sp\|P0AAR5\|YBAN_ECOLI  Structure 020  gene:ybaN  polypeptide:DUF454 domain-containing inner membrane protein YbaN | Hit with:6rfl  z-score:6.8  rmsd:2.8  Aligned residues:57 | Hit with: 7awa  Probability: 88.32  Aligned columns: 34  E-Value: 1.4 | DNA-dependent RNA polymerase  Secretion system apparatus protein  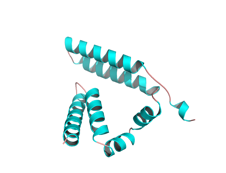 |
| **High confidence**  **Annotation** | b1104 sp\|P75946\|YCFL_ECOLI  Structure 010  gene:ycfL  polypeptide:DUF1425 domain-containing protein YcfL | Hit with:4gio  z-score:12.3  rmsd:1.5  Aligned residues:96 | Hit with: 4gio  Probability: 99.92  Aligned columns:101  E-Value: 2e-23 | Lipoprotein  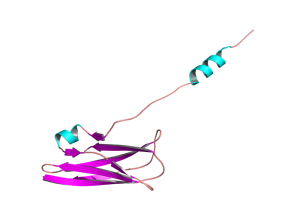 |
| **No Annotation** | Structure 01  b1085 sp\|P62066\|YCEQ_ECOLI  gene:yceQ  polypeptide:DUF2655 domain-containing protein YceQ | Bad model | No hits | =====  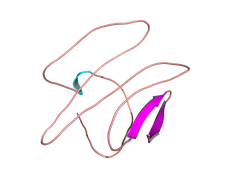 |
| **Low confidence**  **Annotation** | b0232 sp\|Q47156\|YAFN_ECOLI  Structure 017  gene: yafN  polypeptide: antitoxin YafN | Hit with:2a6q  z-score:5.1  rmsd:2.4  Aligned residues:51 | Hit: 2A6Q  Probability: 98.47  Aligned columns: 68  E-Value: 2.3e-6 | Antitoxin  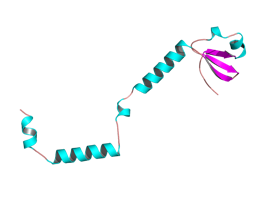 |
| **Low confidence**  **Annotation** | b1848 sp\|P0ACY9\|YEBG_ECOLI  Structure 014  DNA damage-inducible protein YebG | Hit with:4njc  z-score:4.8  rmsd:5.9  Aligned residues:60 | Hit with: PF07130.16  Probability: 100  Aligned columns: 74  E-Value: 1.3e-33 | DNA damage-inducible  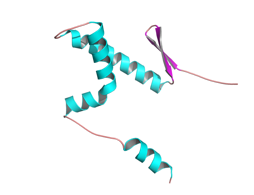 |
| **High confidence**  **Annotation** | b0549 sp\|P68661\|YBCO_ECOLI  Structure 027  gene:ybcO  polypeptide:DLP12 prophage; putative nuclease YbcO | Hit:3g27  z-score:16.7  rmsd:0.7  Aligned residues:81 | Hit: 3g27  Probability: 99.96  Aligned columns:96  E-Value: 1.9e-28 | Prophage  Zinc binding  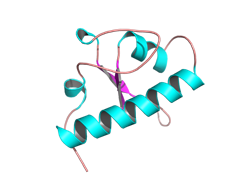 |
| **Low confidence**  **Annotation** | b1777 sp\|P76231\|YEAC_ECOLI  Structure 05  gene:yeaC  polypeptide:DUF1315 domain-containing protein YeaC | Hit:6yt0  z-score:4.9  rmsd:3.3  Aligned residues:53 | Hit: 2hs5  Probability: 91.39  Aligned columns:32  E-Value: 0.37 | Transcriptional regulator  DNA binding  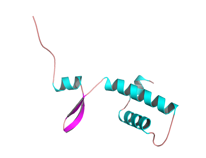 |
| **Low confidence**  **Annotation** | b1332 sp\|P64445\|YNAJ_ECOLI  Structure 025  gene: ynaJ  polypeptide: DUF2534 domain-containing protein YnaJ | Hit:5uz7  z-score:6.6  rmsd:3.1  Aligned residues:69 | Hit: PF10749.13 Protein of unknown function (DUF2534)  Probability: 100  Aligned cols: 80  E-value: 3.4e-34, | Nucleotide Binding Protein  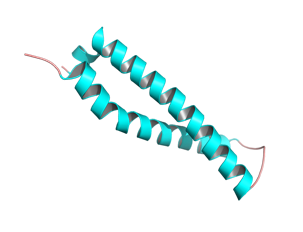 |
| **No Annotation** | b1648 sp\|P64474\|YDHL_ECOLI  Structure 09  gene: ydhL  polypeptide: DUF1289 domain-containing protein YdhL | Bad model | No hits | =====  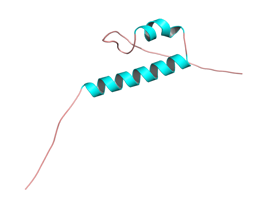 |
| **Low confidence**  **Annotation** | b2377 sp\|P76521\|YFDY_ECOLI  Structure 06  gene:yfdY  polypeptide:DUF2545 domain-containing protein YfdY | Hit:2m6u  z-score:7.2  rmsd:3.4  Aligned residues:74 | Hit: COG2059  Probability: 61.82  Aligned columns:67  E-Value: 100 | Choline Binding Protein  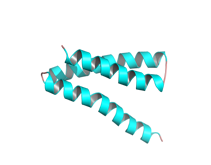 |
| **Low confidence**  **Annotation** | b4406 sp\|P0A8K5\|YAEP_ECOLI  Structure 04  gene:yaeP  polypeptide:PF06786 family protein YaeP | Hit:6kmb  z-score:7.6  rmsd:2.8  Aligned residues:65 | Hit:5h1n  Probability: 100  Aligned columns:66  E-Value: 4.2e-44 | Nuclear Protein  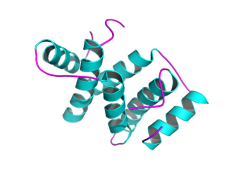 |
| **Low confidence**  **Annotation** | b4176 sp\|P0AF73\|YJET_ECOLI  Structure 021  gene:yjeT  polypeptide:DUF2065 domain-containing protein YjeT | Hit:4xk4-D  z-score: 5.4  rmsd: 4.4  Aligned residues:61 | Hit:3jc8  Probability:60.72  Aligned columns:34  E-Value:69 | HTH transcriptional regulator  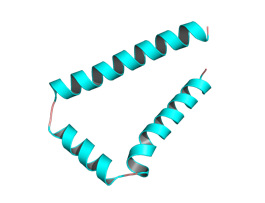 |
| **No Annotation** | b1824 sp\|P64508\|YOBF_ECOLI  Structure 03  gene:yobF  polypeptide:DUF2527 domain-containing protein YobF | Bad model | No hits | =====  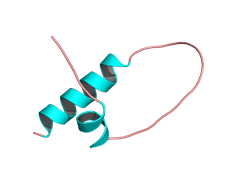 |
| **No Annotation** | b4402 sp\|P0ADD9\|YJJY_ECOLI  Structure 011  gene:yjjY  polypeptide:uncharacterized protein YjjY | Bad model | No hits | =====  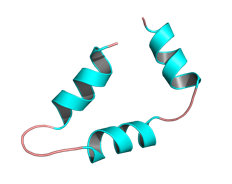 |
